# Genomic and phenotypic consequences of long-term isolation and small population size in Yakushima macaques

**DOI:** 10.64898/2026.10.02.752851

**Authors:** Tsuyoshi Ito, Takashi Hayakawa, Yosuke Kurihara, Goro Hanya, Akihisa Kaneko, Takayoshi Natsume, Seitaro Aisu, Takaaki Honda, Hikaru Wakamori, Shuji Yachimori, Tomoko Anezaki, Toshinori Omi, Shin-ichi Hayama, Yoshi Kawamoto

**Affiliations:** The Kyoto University Museum, Kyoto University, Yoshida Honmachi, Sakyo-ku, Kyoto, Japan; Primate Research Institute, Kyoto University, Kanrin 41-2, Inuyama, Japan; Center for Primate Field Studies, Graduate School of Science, Kyoto University, Sakyo-ku, Kyoto, Japan; Faculty of Environmental Earth Science, Hokkaido University, N10W5, Kita-ku, Sapporo, Japan; Center for Education and Research in Field Sciences, Faculty of Agriculture, Shizuoka University, Nishi-Fujidaira 1623-1, Tenryu-ku, Hamamatsu, Japan; Institute for the Evolutionary Origins of Human Behavior, Kyoto University, Kanrin 41-2, Inuyama, Japan; Faculty of Letters, Research and Development Initiative, Chuo University, 742-1 Higashinakano, Hachioji-shi, Tokyo, Japan; Yokogurayama Natural Forest Museum, Ochihei 737-12, Ochi, Kochi, Japan; Gunma Museum of Natural History, Kamikuroiwa 1674-1, Tomioka, Japan; Faculty of Veterinary Science, Nippon Veterinary and Life Science University, Kyonan-cho 1-7-1, Musashino, Japan; Takagoyama Monkey Observation Club, Horinouchi 2-12-1, Kasugai, Japan

**Keywords:** fluctuating asymmetry, genetic drift, genetic load, inbreeding, island biology, *Macaca*, Yakushima

## Abstract

The relative roles of genetic drift, inbreeding, and natural selection in island evolution have been fundamental themes in evolutionary biology for nearly a century. However, empirical studies have yet to reach a consensus, and research integrating both genotypes and phenotypes in primates remains limited. The present study aims to estimate the demographic history of an island primate population, the Yakushima macaque (*Macaca fuscata yakui*), and evaluate the genomic and morphological consequences of its isolation. To this end, we conducted population genetic analyses based on whole-genome sequencing and geometric morphometric analyses of cranial shape. Our results indicate that *M. f. yakui* has been isolated with an effective population size of several thousand individuals for over 100,000 years. Although we detected signatures of purifying selection, deleterious alleles have not been sufficiently purged. The degree of fluctuating asymmetry in males was slightly, yet significantly, higher in *M. f. yakui* than in mainland populations, whereas this trend was not observed in females. Marginal evidence of directional selection was found for cranial size, but not for shape. Overall, our findings suggest that due to long-term isolation, genetic drift and inbreeding have exerted a substantial influence on shaping genotypic and phenotypic variation in this island population.

---

The evolutionary consequences of population isolation and bottlenecks on islands are fundamental topics in evolutionary biology and conservation ecology. It is intriguing that a bottleneck plays both negative and positive roles in diversification and population fitness. In small populations, genetic drift often overwhelms natural selection; consequently, a decline in population size reduces genetic diversity while increasing inbreeding and realised genetic load. This, in turn, decreases population fitness and elevates the risk of extinction. However, it has been suggested that strongly deleterious mutations are more likely to be purged by purifying selection in smaller populations, as inbreeding exposes these mutations in a homozygous state (Dussex et al. 2023a). Furthermore, genetic drift may enable individuals to cross fitness valleys in the adaptive landscape, facilitating a shift to another adaptive zone that potentially contains higher adaptive peaks (Wright 1932). In larger populations, such valley crossing is unlikely because individuals deviating from local adaptive peaks are more prone to elimination by selection. Therefore, a population bottleneck does not necessarily inhibit diversification or reduce fitness; rather, it may create the potential for future diversification.

Empirical studies regarding the genomic consequences of population bottlenecks have accumulated, yet no consensus has been reached regarding the efficiency of purifying selection. Some researchers argue that strongly deleterious alleles—but not slightly or moderately deleterious ones—are purged in small populations (Xue et al. 2015; Robinson et al. 2018; Grossen et al. 2020; Dussex et al. 2021, 2023b; Khan et al. 2021; von Seth et al. 2021; Wang et al. 2023). In contrast, others have failed to detect efficient purging of strongly deleterious alleles, even in small populations (Wang et al. 2021; Taylor et al. 2024; Bemmels et al. 2025; Gates et al. 2025). Most previous studies have evaluated genomic erosion without investigating its phenotypic effects, with some notable exceptions (Ralls et al. 2000; White et al. 2015; e.g., Huffmeyer et al. 2022). Although directly estimating fitness is challenging, fluctuating asymmetry, a measure of developmental instability, can serve as a proxy for fitness (Møller 1997). Fluctuating asymmetry refers to non-directional deviations from bilateral symmetry (Palmer et al. 1986). Notably, McGrath et al. (2022) reported increased fluctuating asymmetry in mountain gorillas (*Gorilla beringei beringei*), a population with smaller size and lower genetic diversity than other subspecies. Further empirical studies integrating both genotype and phenotype are needed to obtain a comprehensive understanding of the consequences of population bottlenecks.

The roles of genetic drift and natural selection in phenotypic differentiation in island populations have been a subject of controversy for decades. Some empirical studies have found that phenotypic differentiation on islands can be explained primarily by genetic drift (Runemark et al. 2010; Spurgin et al. 2014; Sendell-Price et al. 2020), while others have detected a significant contribution of directional selection (Thorpe and Malhotra 1996; Clegg et al. 2002, 2008; Funk et al. 2016). Although empirical studies testing drift and selection in island phenotypes have accumulated for various taxa, studies targeting non-human primates remain scarce. Addressing this gap offers insights comparable to those gained from humans and other organismal groups.

Here, we investigate the genomic and morphological consequences of long-term isolation and small population size using the Yakushima macaque, *Macaca fuscata yakui* Kuroda, 1940—one of the two subspecies of the Japanese macaque, *M. fuscata* (Gray, 1870)—as a case study. Although *M. fuscata* is a widely used laboratory primate, its evolutionary history and genomic information—particularly that of *M. f. yakui*—remain poorly understood, barring a few studies (Lee et al. 2016; Higashino et al. 2025). *M. f. yakui* is restricted to Yakushima Island (approx. 504 km²), located about 60 km from the mainland, marking the southern limit of the species’ distribution. There has been ongoing debate regarding the phylogenetic relationships and demographic histories of *M. f. yakui* and the mainland subspecies, the Hondo macaque (*M. f. fuscata*). While *M. f. yakui* exhibits distinct morphological traits, including a smaller body, darker pelage, and a narrower face (Kuroda 1940; Yano et al. 2020), some molecular studies have challenged the sister relationship between the two subspecies (Kawamoto et al. 2007; Ito et al. 2021). *M. f. yakui* possesses extremely low genetic diversity, but the timing and extent of its population bottleneck remain unclear, though the Kikai Caldera eruption (ca. 7,300 years ago) is suggested to have had a significant impact (Hayaishi and Kawamoto 2006). By conducting whole-genome and geometric morphometric analyses, this study: (1) estimates the phylogenetic and demographic histories of *M. f. yakui*; (2) evaluates genetic diversity, inbreeding, and genetic load; (3) assesses fluctuating asymmetry as a measure of developmental instability; and (4) tests the neutrality of cranial differentiation. Our data reveal the long-term isolation and small population size of *M. f. yakui*, suggesting that genetic drift and inbreeding have played a dominant role in shaping its genetic and morphological variation.

## MATERIALS & METHODS

### Whole-genome sequencing, genotype calling, annotation, and phasing

We performed whole-genome shotgun sequencing on 17 Japanese macaques (*M. fuscata*) from six populations, including seven individuals from Yakushima (*M. f. yakui*) (Table S1; Fig. 1a). All samples originated from wild individuals, except for two *M. f. yakui* individuals born in captivity and housed at the Primate Research Institute (PRI), Kyoto University. For the Takasakiyama population, we used two blood-derived DNA samples from wild (but provisioned) individuals, stored at the Institute for the Evolutionary Origins of Human Behavior (EHUB), Kyoto University, with permission from Oita City. These samples were originally collected by the former Population Genetics Section of the PRI during a capture survey in 1997. The remaining 15 samples were obtained from previous studies (Suzuki-Hashido et al. 2015; Ito et al. 2021). All samples were collected in compliance with institutional guidelines and approved by relevant authorities; they were obtained during population management surveys or health screenings and reused with appropriate consent.

**Figure 1.**
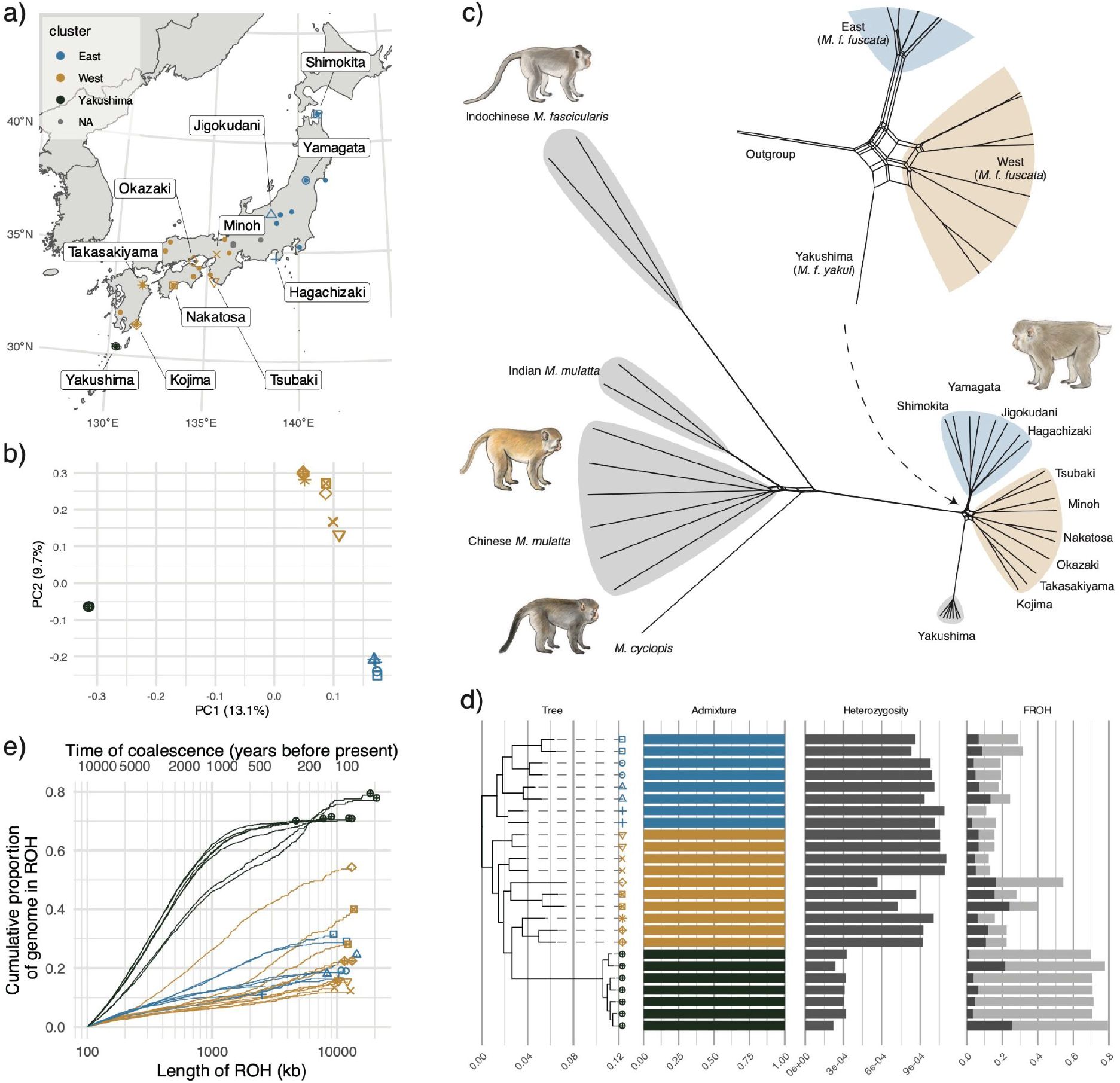
Sampling localities, population structure, and inbreeding. (a) Sampling localities of *M. fuscata*. Large symbols indicate the 11 sampling locations of the DNA samples. Sequence data for the Jigokudani, Hagachizaki, Minoh, Tsubaki, and Okazaki (Shodoshima) populations were obtained from the NCBI SRA (Higashino et al. 2025), while data for the remaining localities were newly sequenced in the present study. Small circles indicate the locations of cranial specimens. Symbol colors correspond to the three major clusters—West (*M. f. fuscata*), East (*M. f. fuscata*), and Yakushima (*M. f. yakui*)—inferred by PCA and ADMIXTURE analyses. Cranial specimens were assigned to clusters based on geographical proximity to the DNA sampling localities, although localities around the putative West–East cluster boundary were left as unassigned. (b) PCA based on autosomal SNVs of all unrelated *M. fuscata* samples. (c) NeighborNet graph of *M. fuscata* and outgroups based on identity-by-state distances. The inset shows a magnified view of the basal reticulate divergence of *M. fuscata*. (d) Population genetic statistics. From left to right: a maximum likelihood tree; genetic clustering inferred from ADMIXTURE at *K* = 3; heterozygosity (per-individual nucleotide diversity); and the ROH-based inbreeding coefficient (*F*_ROH_). *F*_ROH_ represents the sum of the lengths of all ROHs divided by the total length of autosomes. Light gray denotes *F*_ROH≧100kb_, while dark gray denotes *F*_ROH≧2Mb_. (e) Cumulative proportion of the genome in ROH.

Further details regarding these samples are available in the aforementioned publications. Sequencing was performed using 150-bp paired-end reads on Illumina NovaSeq 6000 platform at Macrogen Japan Corp (Tokyo, Japan).

We also incorporated publicly available high-coverage whole-genome shotgun sequences from 14 wild-born Japanese macaques across six populations (Higashino et al. 2025), downloaded from the NCBI Sequence Read Archive (SRA) using SRA Tools (Table S1). Additionally, we included sequences from two Indonesian cynomolgus macaques (*M. fascicularis*), one Taiwanese macaque (*M. cyclopis*), two Indian rhesus macaques (*M. mulatta*), and seven Chinese rhesus macaques as outgroups (Xue et al. 2016; Liu et al. 2018; Bailey et al. 2023; Ray et al. 2025).

Quality control was performed using Fastp v0.24.0 (Chen et al. 2018) with the default settings. The filtered reads were mapped to the NCBI RefSeq assembly of the Indian *M. mulatta* (Mmul_10; GCF_003339765.1) using the “mem” command of Bwa-mem2 v2.2.1 (Vasimuddin et al. 2019). Read group information was added using the “-R” option; for some SRA samples lacking this metadata, the sample ID was added instead. Mapped reads were sorted using Samtools v1.6 (Danecek et al. 2021), and PCR duplicates were marked using MarkDuplicates implemented in the Genome Analysis Toolkit (GATK) v4.6.1.0 (McKenna et al. 2010). Sequencing depth was calculated using Mosdepth v0.3.3 (Pedersen and Quinlan 2018). Parallel processing here and hereafter was performed using GNU Parallel (Tange 2011).

Genotypes were called using GATK HaplotypeCaller with the options “--emit-ref-confidence BP_RESOLUTION --output-mode EMIT_ALL_ACTIVE_SITES”. Joint genotype calling was performed using GATK GenotypeGVCFs with the “--all-sites” option after merging the individual GVCFs via GATK CombineGVCFs. For the X chromosome, this process was conducted separately for each sex. Heterozygous sites are expected to show a unimodal allele fraction distribution peaked at 0.5; however, one sample (#4892) from the Takasakiyama population of *M. f. fuscata* exhibited a trimodal distribution, indicating cross-contamination. Another sample (SRR4454026) from the Henan population of *M. mulatta* showed a high missing rate (>0.4). Joint genotype calling was therefore re-performed excluding these two samples (Table S1), and the resulting dataset was used for downstream analyses. The joint-called VCFs were partitioned into single nucleotide variants (SNVs), short insertions/deletions (indels), and NO_VARIATION using GATK SelectVariants. Hard filtering was applied using GATK VariantFiltration (SNVs: “QD < 2.0, SOR > 3.0, FS > 60.0, MQ < 40.0, MQRankSum < -12.5, ReadPosRankSum < -8.0”; indels: “QD < 2.0, FS > 200.0, SOR > 10.0, ReadPosRankSum < -20.0”). Variants were annotated based on the gene feature format (Mmul_10.105) using SnpEff v5.2 (Cingolani et al. 2012). VCF and BED files were processed using Bedtools v2.31.1 (Quinlan and Hall 2010) and Bcftools v1.19 (Danecek et al. 2021).

Variants were further filtered based on mappability, assessed using GenMap v1.3.0 (Pockrandt et al. 2020) with settings “-K 100 -E 2”. This approach calculates the uniqueness of 100-mers at each genomic position while allowing up to two mismatches. Genomic regions with a mappability score of less than one, or with a depth less than 0.5× or greater than 2.0× the median depth, were masked. Repetitive regions were masked based on pre-existing RepeatMasker results (GCF_003339765.1_Mmul_10_rm.out). For the X chromosome, the pseudoautosomal region (PAR) was also masked. The PAR was identified using a coverage-based approach (Jensen et al. 2024) and subsequently masked. Specifically, the sum of depth in non-overlapping 10-kb windows was calculated separately for both sexes and normalized by the autosomal mean depth. In non-PAR regions, the normalized depth is expected to be twice as high in females as in males, whereas it should be equal in PAR regions. Based on this, the non-PAR region was identified as NC_041774.1:2,350,000–151,960,000.

Read-based phasing for autosomes was performed using WhatsHap v2.6 with default settings (Martin et al. 2016). For the four trios, phasing was conducted by integrating pedigree information alongside read-based data, with the recombination rate parameter set to 0.433 (Xue et al. 2016). Subsequently, population-based phasing for biallelic SNPs was performed using SHAPEIT v4.2.2 (Delaneau et al. 2019) with the options “--effective-size 50000 --use-PS 0.0001 --sequencing”.

Only biallelic variants with a missing rate of less than 0.2 were retained. After all filtering steps, we identified 30,197,539 SNVs and 3,424,558 indels on the autosomes, and 354,426 SNVs on the X chromosome.

### Population genetics

Population genetic analyses were performed using autosomal SNVs. We evaluated relatedness and removed first- and second-degree relatives using PLINK v2.0.0 (Chang et al. 2015; Purcell and Chang 2025) with the “--king-cutoff” parameter set to 0.0884 (Manichaikul et al. 2010). For the *M. fuscata* subset, linkage disequilibrium (LD) was calculated and pruned using Plink v1.9.0 (Gaunt et al. 2007; Purcell and Chang 2024) with the settings “--indep-pairwise 50 5 0.2”. Principal component analysis (PCA) and admixture analysis were performed using PLINK v1.9.0 and ADMIXTURE v1.9.0 (Alexander et al. 2009), respectively. The Tracy-Widom test was conducted using the AssocTests package v1.0.1 (Wang et al. 2020b) in R. The fixation index (*F_ST_*)among the three major clusters (defined below) was calculated following the method of Weir and Cockerham (1984) using PLINK v2.0.0.

For the phylogenetic analyses, distance-based pruning was performed on the dataset including outgroups using the setting “--bp-space 10000”, because correlation-based LD pruning can cause biases in downstream analyses when the dataset contains genetically divergent populations (Bercovich et al. 2025). An identity-by-state (IBS) distance matrix was calculated using PLINK v1.9, and the resulting NeighborNet graph was visualized using the SplitsTree App (Huson and Bryant 2024). Then, a maximum likelihood tree was estimated using IQ-Tree 3 v3.0.1 (Wong et al. 2026). After sites with a zero minor allele count were removed, the variant data were converted to PHYLIP format using vcf2phylip.py (https://github.com/edgardomortiz/vcf2phylip). After removing invariant sites, the substitution model was estimated (Kalyaanamoorthy et al. 2017), and subsequently, the tree was estimated with 100 bootstrap replicates using the settings “-m MFP+ASC --mrate E,G -b100”. The TVM+F model was selected.

Population-level maximum likelihood trees incorporating potential migration events were constructed using TreeMix v1.13 (Pickrell and Pritchard 2012). Using the distance-pruned dataset, trees were generated with Indochinese *M. fascicularis* as the outgroup and 0–5 migration edges using the parameters “-k50 -noss”.

Gene flow between populations was estimated based on the unpruned SNVs using Dsuite v0.5 (Malinsky et al. 2021). The *D* and *f*_4_-ratio statistics were calculated for all possible population trios using the “Dtrios” or “DtriosParallel” commands. We subsequently calculated and visualized *f*-branch statistics (Malinsky et al. 2018) based on a TreeMix tree with zero migration edges using the “Fbranch” command and dtools.py (https://github.com/millanek/Dsuite) with some modifications. In these analyses, two Indonesian *M. fascicularis* served as the outgroup. To evaluate sex-biased gene flow, we compared *f*-branch statistics between the autosomes and the non-PAR X chromosome.

Admixture history was further investigated based on the unpruned SNVs using ADMIXTOOLS 2 v2.0.10 (Maier et al. 2023). Outgroup *f*_3_ statistics were calculated to estimate genetic similarities between *M. fuscata* populations and other species, using *M. fascicularis* as an outgroup. A qpWave analysis was performed to evaluate the minimum number of migration waves from the source populations to *M. fuscata*. Focusing specifically on *M. f. yakui*, we calculated outgroup *f*_3_-statistics to estimate genetic similarities with *M. f. fuscata* populations, alongside *f*_4_ statistics in the form of *f*_4_(Indochinese *M. fascicularis*, *M. f. yakui*; X, Shimokita *M. f. fuscata*) to test for gene flow. The *f*_4_ statistics were compared between autosomes and the non-PAR X chromosome. Finally, admixture graphs with 0–5 migration edges were fitted using Indochinese *M. fascicularis* as the outgroup, excluding populations represented by fewer than two samples.

Demographic history was inferred from the phased SNVs using MSMC2 v2.1.4 (Wang et al. 2020a). Two samples per population were included; the Okazaki (Shodoshima) and Takasakiyama populations were excluded as they contained only one unrelated individual. Additionally, an analysis using five wild individuals of *M. f. yakui* was performed to focus on the more recent past. Input files were generated using generate_multihetsep.py from the msmc-tools repository (https://github.com/stschiff/msmc-tools). MSMC2 was run with time segment settings of “30*1+1*2+1*4+1*6”, which was adjusted to “35*1+1*2+1*4+1*6” for the five-sample *M. f. yakui* dataset. To estimate population divergence times, the relative cross-coalescent rate (rCCR) was calculated using combineCrossCoal.py. To validate recent population size fluctuations in *M. f. yakui*, SMC++ v1.15.4 (Terhorst et al. 2017) was applied to all seven individuals with the options “--timepoints 10 10000 --regularization-penalty 4”. We assumed a mutation rate of 7. 7 × 10 per site per generation and a generation time of 10.4 years (Bergeron et al. 2021).

The demographic trajectory of *M. f. yakui* was further evaluated using Fastsimcoal2 v2.8 (Excoffier et al. 2021). We tested four hypothesized models (Models 0–3). Guided by the MSMC2 results, all models assumed an ancestral reduction in population size in the distant past. Model 0 assumed a constant population size thereafter; Model 1 assumed a recent expansion; Model 2 assumed a recent decline; and Model 3 assumed a bottleneck followed by recovery to pre-bottleneck levels. To identify derived alleles, we retained only sites with an alternative allele frequency < 0.2 across the outgroups (*M. fascicularis*, *M. cyclopis*, and *M. mulatta*). The unfolded site frequency spectrum was constructed from autosomal, biallelic derived SNVs in intergenic regions using easySFS.py (https://github.com/isaacovercast/easySFS). One hundred independent Fastsimcoal2 runs were performed for each model, with 50 expectation-conditional maximization (ECM) cycles (“-L 50”) and 200,000 simulations (“-n 200,000”). Model selection was based on the Akaike Information Criterion (AIC). To validate the selection, we evaluated the likelihood distribution by running the best-fitting model 100 times with 10 million simulations, following Meier et al. (2017). Confidence intervals for parameter estimates were obtained via a non-parametric bootstrap approach with 100 replicates, using genomic data split into 100 blocks.

To assess inbreeding, runs of homozygosity (ROH) were identified from autosomal SNVs (without LD pruning) using PLINK v1.9.0 with settings “—homozyg --homozyg-density 1000 --homozyg-kb 100”. The ROH-based inbreeding coefficient (*F*_ROH_) was calculated as the cumulative length of ROHs divided by the total autosomal length. The timing of ROH occurrence was estimated using the formula *g* = 100/(2*rL*), where *g* is the number of generations back in time, *r* is the recombination rate, and *L* is the ROH length in Mb (Kyriazis et al. 2025). The recombination rate was assumed to be 0.433 cM/Mb (Xue et al. 2016), with a generation time of 10.4 years (Bergeron et al. 2021). Per-sample nucleotide diversity (heterozygosity) was calculated using Pixy v1.2.10 (Korunes and Samuk 2021).

To evaluate genetic load, the derived variants were categorized into loss-of-function (LoF), missense, synonymous, and intergenic classes based on SnpEff annotations. For each mutation category, we calculated the number of homozygous derived sites (realized load), sites with at least one derived allele (potential load), and the total number of derived alleles (total load) per individual. To account for varying missing data rates, counts were normalized by the number of callable sites per individual, following Marsden et al. (2016). Specifically, raw counts were divided by the individual’s callable sites and multiplied by the mean number of callable sites across all individuals. Finally, the nonsynonymous-to-synonymous variant ratio, based on missense mutations, was calculated for heterozygous sites.

Genetic load was also evaluated at the population level. Based on the PCA and admixture results, three clusters were defined: East, West, and Yakushima. We calculated *R_XY_* (Xue et al. 2015) as the relative frequency of derived alleles in one cluster compared to another for each mutation category. The *R_XY_* of each functional category was normalized by the *R_XY_* of intergenic regions. Standard errors were estimated using a 100-block jackknife procedure. Finally, we visualized the site frequency spectrum of polymorphic sites for each mutation category and cluster. To correct for sample size differences across clusters, we downsampled the data to 14 alleles using a hypergeometric distribution over 100 replicates. The mean and 95% confidence intervals across replicates were visualized.

### Geometric morphometrics

We analyzed 331 crania from 22 populations of *M. fuscata*, 68 of which were from *M. f. yakui* (Table S2; Fig. 1a). The specimens are housed at the Gunma Museum of Natural History (Tomioka, Japan), the Shikoku Institute of Natural History (Nakatosa, Japan), the Hakusan Nature Conservation Center (Hakusan, Japan), the Tochigi Prefectural Museum (Utsunomiya, Japan), and the EHUB, to which the collection from the former PRI was transferred. The individuals and populations examined here do not necessarily correspond to those used for DNA analysis. All crania were from wild adults with fully erupted third molars.

Crania were scanned at the PRI using an Asteion Premium 4 helical scanner (Toshiba Medical Systems, Otawara, Japan) with a slice thickness of 0.5 mm. DICOM slices were reconstructed at 0.2 mm intervals. Images were segmented using edge detection and the watershed method (Ito 2019), and three-dimensional (3D) surface models were generated using create_mesh.py, a script based on dicom2stl (https://github.com/itots/dicom2stl).

Forty-five anatomically defined 3D landmarks (Table S3; Fig. S1) were digitized on the surface models using 3D Slicer 5 (Kikinis et al. 2014; https://www.slicer.org/). To assess and mitigate measurement error, a single author (T.I.) repeated the digitization four times for all specimens. For two of the four replicates, digitization was performed on mirrored surface models to account for potential handedness bias; coordinates were subsequently re-mirrored for analysis.

Missing landmarks (shown in 96 specimens; range 1–8, mean 1.9) were estimated using Bayesian PCA via the “MissingGeoMorph” function of the LOST package v2.1.3 (Arbour and Brown 2014, 2025) in R, separately for each replicate. Given the sexual dimorphism in macaque crania, subsequent analyses were conducted separately for each sex unless otherwise stated. Measurement error was evaluated, and generalized Procrustes Analysis (GPA) was performed using the “bilat.symmetry” function of the geomorph package v4.1.0 (Baken et al. 2021; Adams et al. 2025) in R. Measurement errors were deemed acceptable relative to individual differences and fluctuating asymmetry (Table S4). The symmetric shape component was used for subsequent analyses unless otherwise stated. Centroid size was calculated as the square root of the sum of squared distances of all landmarks from their centroid, averaged across the four replicates. PCA was performed on the symmetric and asymmetric shape components using the “gm.prcomp” function of geomorph. Allometry was assessed using the “procD.lm” and “plotAllometry” functions, with shape as the response variable and centroid size and subspecies as explanatory variables. Interspecific shape differences were further evaluated based on between-group PCA using “groupPCA” function in the Morpho package v2.13 (Schlager 2017). Shape changes were visualized via surface models deformed by thin-plate splines, implemented in the Morpho package.

Fluctuating asymmetry was quantified using an unsigned asymmetry index (Lazić et al. 2015) derived from the bilat.symmetry results. This index was scaled by subtracting one and multiplying by 1,000. Bayesian regression was performed in R using the brms package v2.23.0 (Bürkner 2017), with the scaled unsigned asymmetry index as the response variable, subspecies as a fixed effect, and population as a random effect. Populations with fewer than four individuals were excluded from the analysis. We assumed a lognormal distribution and specified weakly informative priors using a Student’s *t*-distribution, *t*(3, 0, 10), truncated at zero for standard deviation parameters. We also analyzed sex-combined data, including subspecies, sex, and the sex–subspecies interaction as fixed effects. To control for potential bias from missing landmarks or qualitative anomalies (e.g., tooth agenesis, supernumerary or rotated teeth, partial bone swelling/depression, deformation), analyses were replicated using a subset of 228 intact specimens.

We tested the evolutionary process of cranial diversification among populations using the beta test (Ackermann and Cheverud 2002, 2004), based on Lande’s quantitative genetic model (Lande 1980). Neutral diversification via genetic drift follows the equation ***B*** = (*t*/*N_e_*)***G***, where ***B*** is the between-group variance/covariance (V/CV) matrix, ***G*** is the additive genetic V/CV matrix of the founding population, and *t* and *N_e_* are the number of generations and effective population size, respectively. Since ***G*** is often assumed to be proportional to the within-group phenotypic V/CV matrix (***W***) (Cheverud 1988), and *t*/*N_e_* is constant for specific comparisons, ***B*** is expected to be proportional to ***W***. The proportionality (***B*** ∝ ***W***) was tested using linear regression of log-transformed between-group variances against the eigenvalues of ***W***. ***W*** was calculated using the “vcvComp” package v1.0.2 (Le Maître and Mitteroecker 2019) as the pooled within-group V/CV matrix of populations with *N* > 20 (Shimokita, Tone, and Yakushima). Between-group variances were calculated using PCs of group means based on ***W***, excluding populations with fewer than four individuals. For the slope test, we retained the first PCs accounting for >90% of cumulative variance (22 for females, 23 for males) to avoid redundancy, following Baab (2021). Using the car package v3.1.5 (Fox and Weisberg 2019), we tested whether the regression slope deviated significantly from one. Significant deviation implies that diversification cannot be explained by drift alone.

We then applied the multivariate drift test (Hohenlohe and Arnold 2008) using the "MultivDriftTest" function in the evolqg package v0.3.6 (Melo et al. 2015) to test whether the magnitude of divergence of *M. f. yakui* from *M. f. fuscata* exceeds neutral expectations. This function simulates population evolution via drift starting from a reference population (*M. f. fuscata*) under the model ***B=(***t/N_e_)**W b** is the expected between-group V/CV matrix. ***W*** was calculated using the “vcvComp” package as the pooled within-group V/CV matrix of populations with *N* > 20 (Shimokita, Tone, and Yakushima). Here, we assumed that the proportion of phenotypic variance due to additive genetic effects is the same for between-population variance and within-population variance. Based on MSMC2 results, we set *t* = 22, 437, which corresponds to the time where rCCR dropped below 0.5 in all population pairs between *M. f. yakui* and *M. f. fuscata*, and N_e_=7,893, which corresponds to the time-weighted harmonic mean of the N_e_ trajectory from the assumed divergence time up to the most recent reliable interval (i.e., excluding the first two intervals). We used the first 25 PCs (>90% variance) for both sexes. Note that the PCs used here were based on the V/CV of the total sample, distinct from the previous beta test. We performed 10,000 simulations and calculated the Euclidean norms of divergence from *M. f. fuscata*. If the observed norm was significantly larger or smaller than expected under the simulated distribution, we rejected the null hypothesis of drift, indicating directional selection or stabilizing selection, respectively.

Finally, we performed a *P_ST_- F_ST_* comparison to test the neutrality of morphological differentiation of *M. f. yakui*. Phenotypic divergence (*P_ST_*) is defined as:

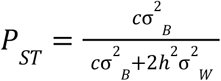

and can be rewritten as:

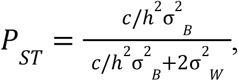

Where σ^2^*_B_* is the between-population phenotypic variance, σ^2^*_W_* is the within-population phenotypic variance, ℎ^2^ is the narrow-sense heritability, and *c* is the scalar describing the proportion of total variance due to additive genetic effects across populations (Brommer 2011). We utilized variance components instead of mean squares for σ^2^*_B_* to rigorously exclude error variance. Consequently, our *P_ST_* estimates are more conservative than those calculated using the widely used Pstat package (Da Silva and Da Silva 2018). Variance components were calculated using the VCA package v1.5.2. We calculated *P_ST_* for centroid size and the first four PCs between *M. f. yakui* and the West clade. Confidence intervals were estimated from 1,000 bootstrap replicates. While the null assumption is ℎ^2^ = *c*, their actual values are unknown; thus, we calculated *P_ST_* varying the ratio *c*/ℎ^2^from zero to two in 0.1 increments. The *F_ST_* value used was a ratio of averages based on the method of Weir and Cockerham (1984), which is appropriate for comparison with *P_ST_* estimates derived using Bessel’s correction (Liu and Edge 2025). If *P_ST_* is significantly larger or smaller than *P_ST_*, the phenotypic divergence cannot be attributed to genetic drift alone, indicating directional or stabilizing selection, respectively.

## RESULTS

### Population structure and phylogeny

PCA identified three clusters within *M. fuscata*: East (Shimokita, Yamagata, Jigokudani, and Hagachizaki populations of *M. f. fuscata*), West (Minoh, Okazaki, Tsubaki, Nakatosa, Takasakiyama, and Kojima populations of *M. f. fuscata*), and Yakushima (*M. f. yakui*) (Fig. 1b). The Tracy-Widom test detected two significant eigenvalues at a significance level of 0.05. PC1 separated *M. f. yakui* from the mainland populations, while PC2 captured the East–West divergence. Admixture analysis supported the PCA findings. The cross-validation error was lowest at *K* = 2 (0.535), separating *M. f. yakui* from the others (Fig. S2). The error was the second lowest by a narrow margin at *K* = 3 (0.548), where the East, West, and Yakushima clusters were detected (Fig. 1d). No indication of genetic admixture was observed between the clusters. The *F_ST_* values were 0.344 between the West cluster and *M. f. yakui*, 0.446 between the East cluster and *M. f. yakui*, and 0.124 between the West and East clusters.

The NeighborNet graph showed that the basal divergence pattern of *M. fuscata* was star-like and reticulated (Fig. 1c). The maximum likelihood tree indicated that *M. fuscata* formed a monophyletic clade, sister to a clade composed of *M. cyclopis*, Chinese *M. mulatta*, and Indian *M. mulatta* (Figs. 1d and S3). *M. f. yakui* placed within the West cluster of *M. f. fuscata*, showing sister-relationship with the Kyushu clade (Takasakiyama and Kojima populations).

### Genetic diversity and inbreeding

Autosome-wide heterozygosity (per-individual nucleotide diversity) was significantly lower in *M. f. yakui* than in *M. f. fuscata* (*t* = 18.09, *df* = 22.40, *p* = 7.4 × 10^−15^), averaging 0.0291% (range: 0.0223–0.0324%) (Fig. 1d). Even after excluding the two captive individuals, the average remained 0.0295% (0.0223–0.0318%). The Okazaki (Shodoshima) population in *M. f. fuscata* exhibited the second-lowest heterozygosity, averaging 0.0567%. Other *M. f. fuscata* populations showed relatively uniform heterozygosity values of approximately 0.1%, although the Nakatosa and Shimokita populations exhibited slightly lower values around 0.08%.

The inbreeding coefficient (*F*_ROH≧100kb_) exhibited an inverse relationship with heterozygosity (Fig. 1d). *M. f. yakui* showed significantly higher *F*_ROH≧100kb_ values (70.1–79.7%) compared to the relatively low values of around 20% in other *M. f. fuscata* populations (*t* = −17.04, *df* = 22.97, *p* = 1.6 × 10^−14^). However, when excluding short ROHs (ROH < 2Mb), *F*_ROH≧2Mb_ did not differ significantly between *M. f. yakui* and *M. f. fuscata* populations (*t* = −0.23, *df* = 7.70, *p* = 0.823). In *M. f. yakui*, ROH lengths were mostly shorter than 1–2 Mb, corresponding to ROH coalescence at 0.6–1.2 ka or older (Fig. 1e). The Okazaki (Shodoshima) population exhibited the second-highest *F*_ROH≧100kb_ (0.543).

The two captive individuals did not differ significantly from other *M. f. yakui* individuals in heterozygosity (*t* = −0.30, *df* = 1.37, *p* = 0.803) or *F*_ROH≧100kb_ (*t* = 0.31, *df* = 1.40, *p* = 0.799). Thus, these individuals were included in subsequent analyses unless otherwise stated.

### Gene introgression

The population-level maximum likelihood tree generated by TreeMix without migration edges placed *M. f. yakui* outside the West clade (Fig. S4). However, allowing a single migration edge revealed gene flow from the *M. f. yakui* lineage to the Kyushu clade (Takasakiyama and Kojima). When 2–5 migration edges were modeled, *M. f. yakui* formed a sister relationship with *M. f. fuscata*, and a strong migration edge was detected from the ancestral *M. fuscata* lineage to that of the East clade.

The *f*-branch statistics indicated that *M. cyclopis* and the ancestral lineage of Chinese *M. mulatta* (and some of its descendants) shared excess alleles with *M. fuscata* (Fig. 2a). The extent of excess allele sharing with *M. mulatta* or *M. cyclopis* was relatively uniform across *M. fuscata* populations (Figs. 2a, b). Notably, the *f*-branch statistics between the ancestral lineage of Chinese *M. mulatta* and *M. fuscata* were substantially lower on autosomes (0.0050–0.0065) than on the non-PAR X chromosome (0.0240–0.0309), supporting the findings of Osada et al. (2021). Within *M. fuscata*, significant excess allele sharing was detected in many population pairs, even across the three major clades. *M. f. yakui* shared excess alleles with some West clade lineages; however, no excess sharing was detected with the East clade, with the exception of the Jigokudani population (Figs. 2a, c). The *f*-branch statistics for the West clade lineages were markedly lower on autosomes than on the non-PAR X chromosome.

**Figure 2.**
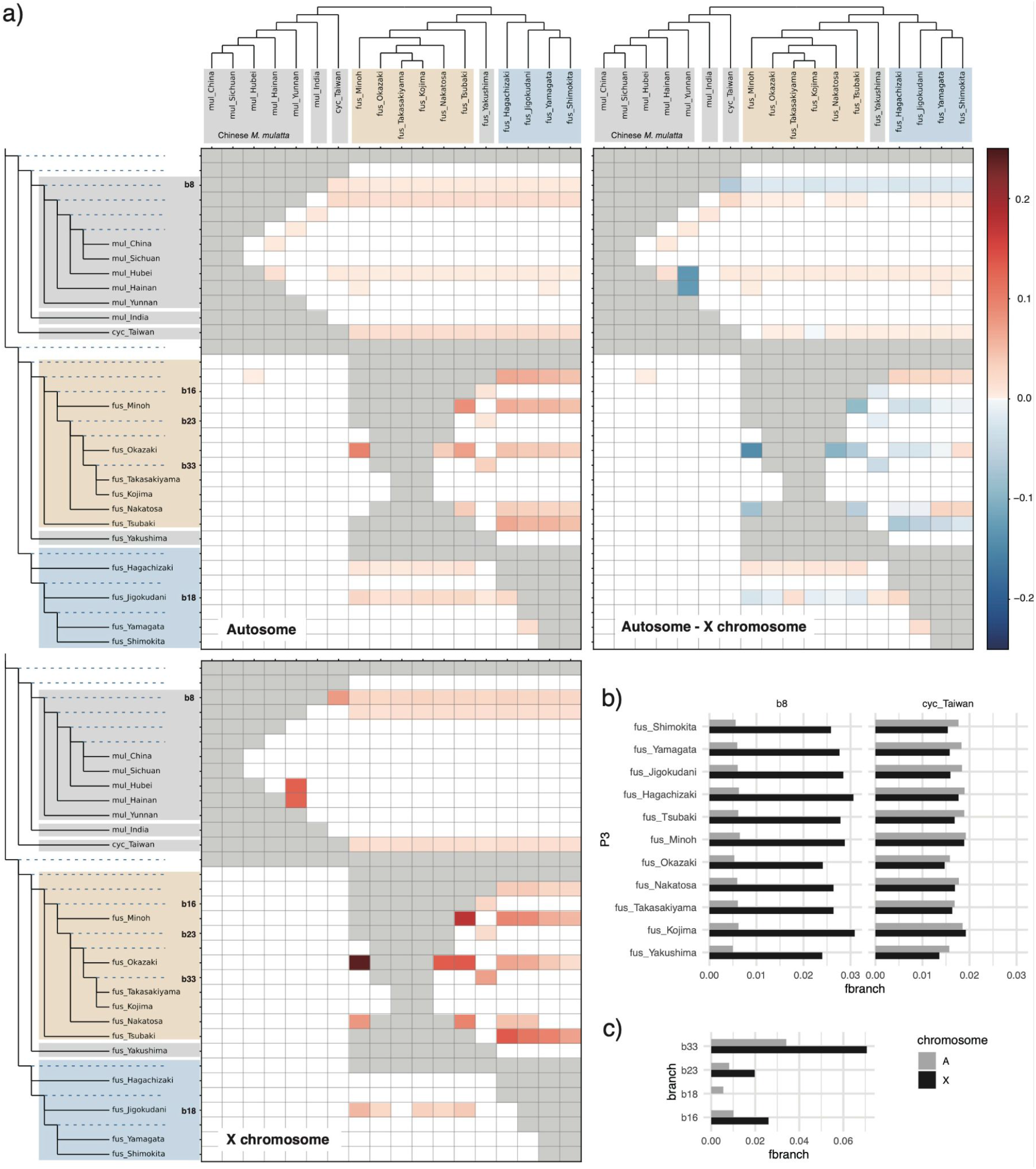
Excess allele sharing between branches and populations. (a) Results of *f*-branch statistics for autosomes (top left) and the non-PAR X chromosome (bottom left). The matrix values (*f*-branch statistics) quantify the degree of excess allele sharing between the branch on the *y*-axis and population P3 on the *x*-axis. The matrix in the top right panel shows the difference between the *f*-statistics of the non-PAR X chromosome and the autosomes; positive values indicate greater excess allele sharing in autosomes compared to the non-PAR X chromosome. Abbreviations: fus, *M. fuscata*; cyc, *M. cyclopis*; and mul, *M. mulatta*. (b) *f*-branch statistics between the ancestral branch of Chinese *M. mulatta* (b8) (left) or *M. cyclopis* (right), and *M. fuscata* populations (P3). (c) *f*-branch statistics measuring excess allele sharing between specific branches of *M. f. fuscata* and the Yakushima population (*M. f. yakui*) as P3. Branch combinations with zero f-branch statistics or those unavailable due to tree topology are omitted.

The qpWave analysis indicated that a single migration wave was sufficient to explain the formation of *M. fuscata* (Table S5). Outgroup *f*_3_ statistics showed that genetic similarity to *M. fuscata* was higher in *M. cyclopis*, followed by the Hainan, Hubei, Sichuan, Yunnan, and Indian populations of *M. mulatta*, in that order; in contrast, these similarities were homogeneous across *M. fuscata* populations (Fig. S5). For *M. f. yakui*, outgroup *f*_3_ statistics indicated that genetic similarity to other *M. fuscata* populations was highest in the Kyushu populations (Kojima and Takasakiyama), followed by the Shikoku–Kinki–Shodoshima populations (Nakatosa, Okazaki, Minoh, and Tsubaki) and the Chubu–Tohoku populations (Hagachizaki, Jigokudani, Yamagata, and Shimokita), in that order (Fig. S6). The *f*_4_ statistics followed this trend but were markedly lower on the non-PAR X chromosome than on autosomes, supporting the *f*-branch results (Fig. S7). In the admixture graph analysis, the worst residual decreased markedly from zero to five migration edges (Table S6), indicating that multiple migration events occurred. Admixture graphs with 3–5 migration edges indicated that *M. f. yakui* has phylogenetic closeness to the West clade populations, but multiple migration events occurred from the East clade populations to the West clade populations; the proportion of migration was lower in the Kyushu (Kojima) population than in the others (Fig. S8).

### Demography and divergence

MSMC2 analysis indicated that the ancestral lineage of *M. fuscata* experienced a strong population bottleneck around 400–500 ka (Fig. 3a). The effective population size continued to decrease in *M. f. yakui*, whereas it recovered to a peak around 100 ka in other populations. Subsequently, many populations, including *M. f. yakui*, experienced a second bottleneck around 10–20 ka. The population size of *M. f. yakui* reached a minor peak around 2–3 ka and then decreased again. SMC++ analysis also identified a weak bottleneck around 10–20 ka, followed by a slight recovery (Fig. 3b).

**Figure 3.**
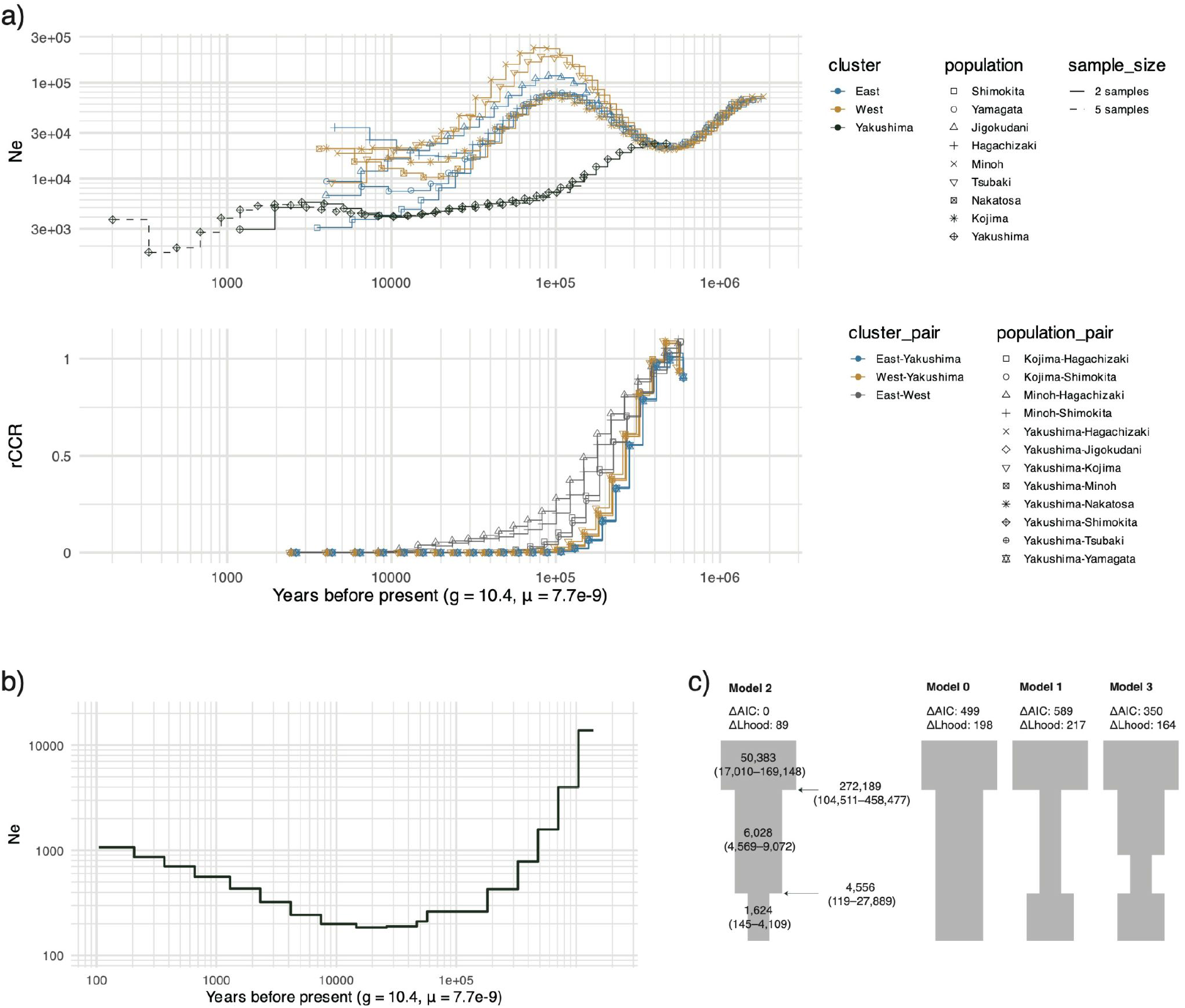
Demographic history. (a) Effective population size over time (top) and rCCR (bottom) inferred using MSMC2. The first two and last six time segments were excluded. (b) Effective population size over time for the Yakushima population (*M. f. yakui*) inferred using SMC++. The oldest time segment and segments corresponding to ≤ 10 years were excluded. (c) Demographic models for *M. f. yakui* inferred using fastsimcoal2. Parameter estimates and their 95% confidence intervals (CIs) are shown for the best-fitting model (Model 2). Effective population sizes are expressed as the number of diploid individuals, and time is given in years.

The rCCR indicated that divergence between the West and East clade populations occurred around 150–200 ka (Fig. 3a). The divergence of *M. f. yakui* from the West and East clade populations was estimated at around 200–300 ka. The gradual decline in rCCR indicated that the divergence of *M. f. yakui* from *M. f. fuscata* was characterized by continuous gene flow during the initial stages, followed by restricted gene flow after approximately 100 ka.

For the fastsimcoal2 simulation, Model 2 (assuming a recent decline) was selected based on the AIC (Fig. 3c). The likelihood distribution of this model did not overlap with that of other models, confirming the robustness of this selection (Fig. S9). According to this simulation, *M. f. yakui* experienced a reduction in effective population size from 6,028 (95% CI: 4,569–9,072) to 1,624 (95% CI: 145–4,109) at 4,556 years ago (95% CI: 119–27,889).

### Genetic load

The number of homozygous derived sites (realized load) showed a positive correlation with *F*_ROH≧100kb_ for LoF, missense, and synonymous variants (Fig. 4d–f). The number of derived alleles (total load) remained constant across individuals for synonymous sites (Fig. 4i; *R*^2^ = 0.00, *p* = 0.99). In contrast, LoF and missense sites exhibited a quadratic relationship with inbreeding levels: the derived allele count decreased with increasing *F*_ROH≧100kb_ within *M. f. fuscata*, whereas it was substantially higher in *M. f. yakui* despite its much higher *F*_ROH≧100kb_ values (Fig. 4g, h). The number of sites carrying derived alleles (potential load) generally showed a negative relationship with *F*_ROH≧100kb_, but this trend for LoF and missense sites plateaued in *M. f. yakui* (Fig. 4a–c). The LoF of indels showed a pattern similar to that of SNVs (Fig. 4j, k, l). The nonsynonymous-to-synonymous ratio for missense mutations was much higher in *M. f. yakui* than in the *M. f. fuscata* populations (Fig. 4m). *R_XY_* statistics indicated that derived alleles of LoF and missense SNVs were more enriched in *M. f. yakui* than in the East or West clades, although the 95% confidence interval included one for the East–*M. f. yakui* comparison for LoF variants (Fig. 5a). Conversely, synonymous SNVs were accumulated slightly more frequently in the East and West clusters than in *M. f. yakui*. Differences between the East and West clades were relatively minor, with confidence intervals including one for all mutation categories.

**Figure 4.**
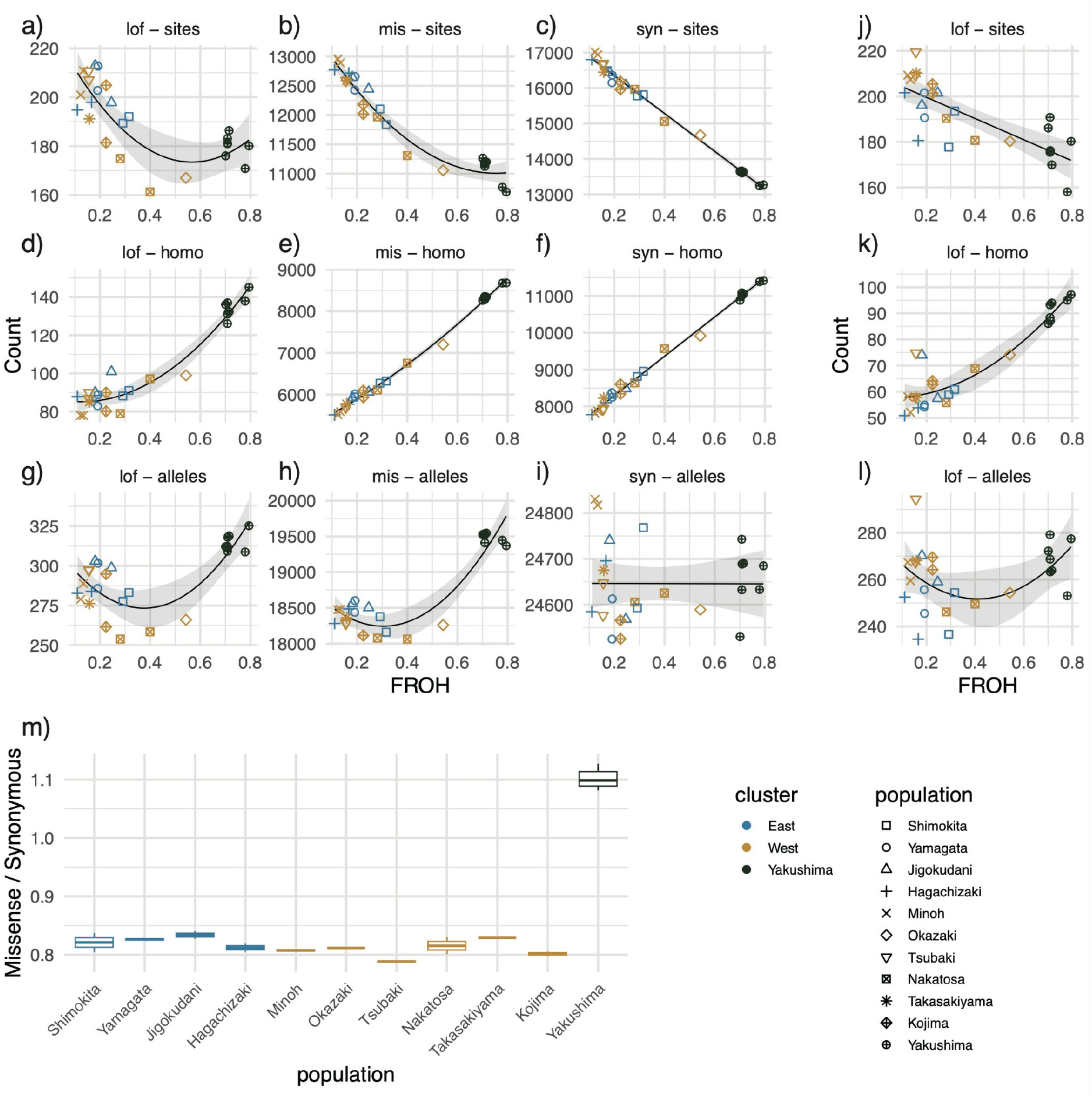
Individual-level genetic load. (a–i) SNVs. (j–l) indels. The *x*-axis represents the ROH-based inbreeding coefficient (*F*_ROH≧100kb_). The *y*-axis rows correspond to the number of sites with at least one derived allele (potential load; a–c, j), the number of homozygous derived sites (realized load; d–f, k), and the total number of derived alleles (total load; g–i, l). Regression lines (linear or quadratic, selected based on the Bayesian information criterion [BIC]) are displayed with 95% confidence intervals. (m) Autosome-wide nonsynonymous-to-synonymous ratio for missense mutations at heterozygous sites.

**Figure 5.**
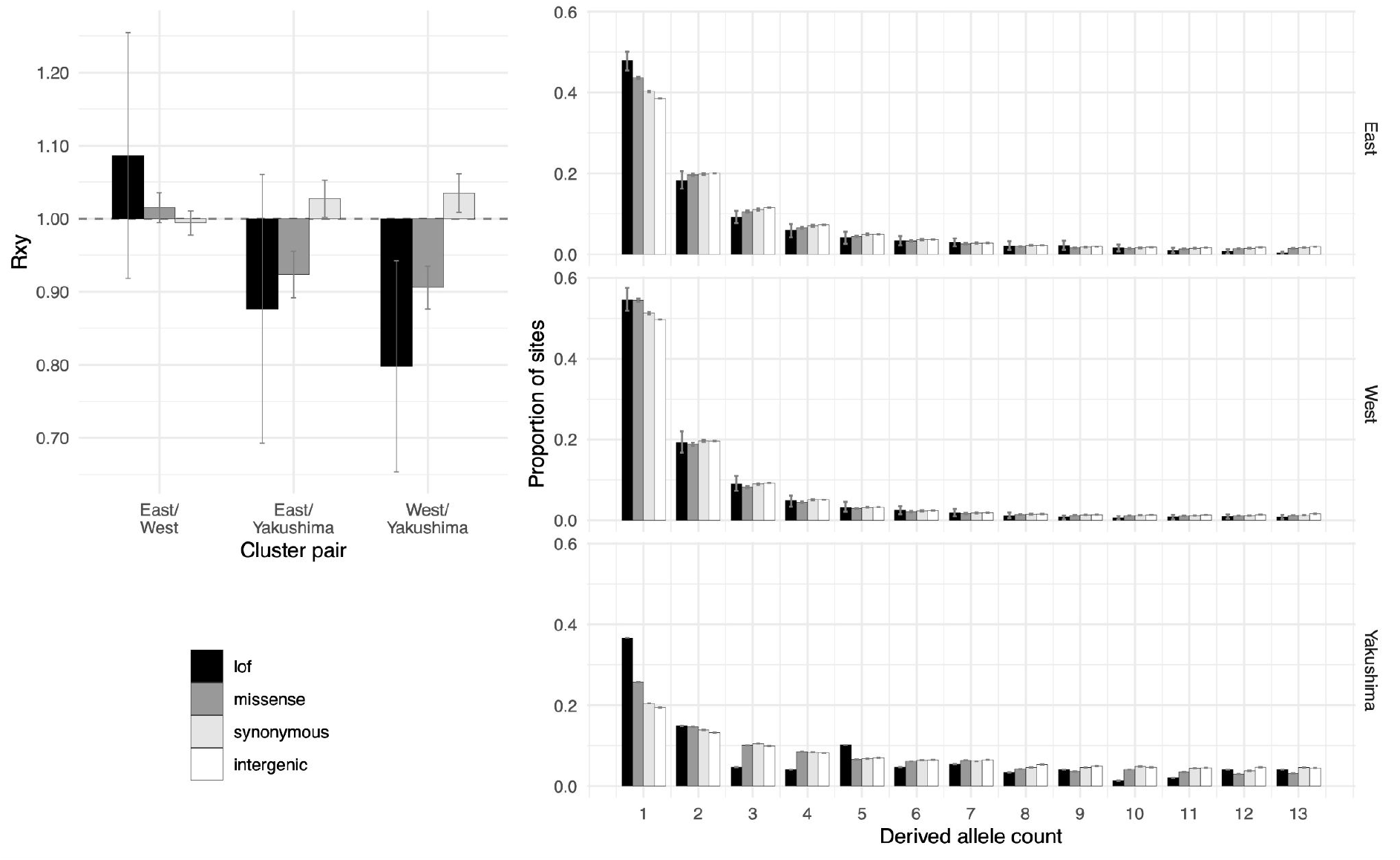
Population-level genetic load. (a) Frequency of derived alleles in population X relative to population Y (*R*_XY_). Error bars represent ±1.96 standard errors. (b) Site frequency spectrum of polymorphic sites. Error bars represent 95% confidence intervals.

In general, the site frequency spectra of the West and East clades were skewed toward lower frequency variants, particularly singletons, compared to *M. f. yakui* (Fig. 5b). When comparing mutation categories within singletons, the relative frequency was highest for LoF variants, followed by missense, synonymous, and intergenic variants. The disparity across mutation categories at singletons was most pronounced in *M. f. yakui*. Beyond singletons, the frequency of LoF variants was lower than that of other mutation categories, a trend that was particularly evident in *M. f. yakui*.

### Morphological variation

PCA demonstrated that the two subspecies were differentiated in cranial shape, although their distributions overlapped considerably along the first two PCs (Fig. 6a). Subspecies differences were clearly evident in the allometry plot. *M. f. yakui* was significantly smaller in centroid size than *M. f. fuscata*. In both subspecies, larger individuals tended to exhibit a more ventrally oriented rostrum, associated with a longer facial profile and a relatively narrower neurocranium (Figs. 6c and S10b). *M. f. yakui* significantly deviated from this common allometric trend (Table S7), exhibiting a peramorphic cranial shape despite its smaller size. bgPCA further resolved the subspecies differences (Figs. 6b and S10a). *M. f. yakui* was characterized by a narrower face and narrower neurocranium, and a more superiorly positioned zygomatic arch compared to *M. f. fuscata* (Fig. 6d).

**Figure 6.**
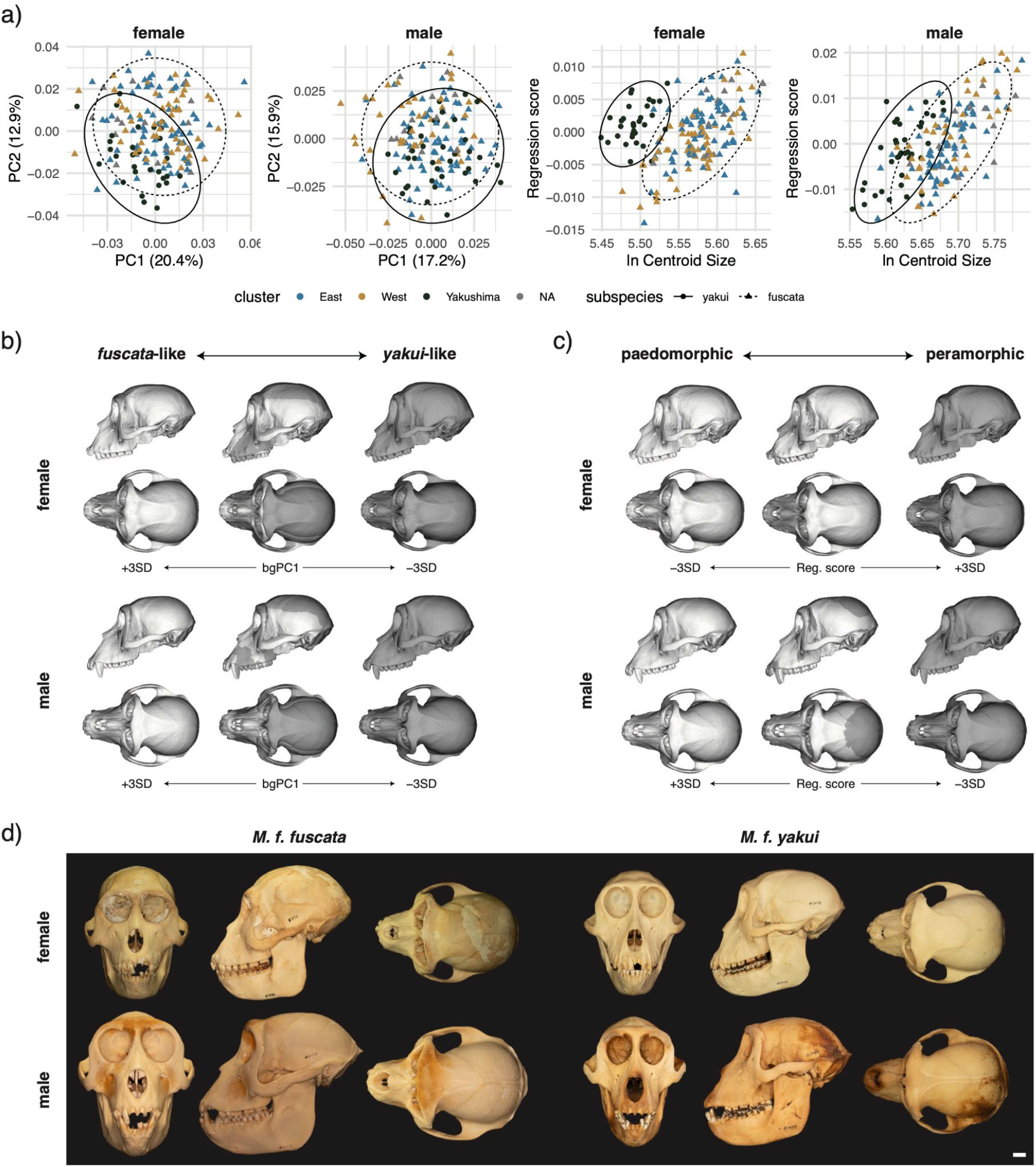
Cranial symmetric shape variation in *M. fuscata*. (a) Scatter plots of the first two PCs (left two panels) and allometric regression scores against ln centroid size (right two panels) for females and males. Subspecies distributions are enclosed by 95% confidence ellipses. (b) Inter-subspecies shape differences analyzed via between-group PCA. (c) Allometric shape changes associated with regression scores. The left and right models represent extreme shape configurations (±3 SD), while the center shows their superimposition. Reference meshes: GMNH-VM10-348 (female) and GMNH-V14-611 (male). See Figure S10 for vector-based shape visualizations. (d) Representative cranial specimens: female *M. f. fuscata* (PRI-5736, upper left), male *M. f. fuscata* (PRI-4113, lower left), female *M. f. yakui* (PRI-3448, upper right), and male *M. f. yakui* (PRI-7999, lower right). Backgrounds were masked using Adobe Photoshop. Scale bar indicates centimeters.

### Fluctuating asymmetry as a measure of developmental instability

Bayesian regression analysis indicated that the scaled asymmetry index was higher in *M. f. yakui* than in *M. f. fuscata* for males (subspecies effect: 0.40 [95% CrI 0.05, 0.75]), but no significant difference was detected for females (0.22 [−0.16, 0.62]) (Table 1; Fig. 7). This pattern remained consistent even when excluding individuals with missing landmarks or qualitative anomalies (Table S8; Fig. S11). Even among males, the distributions overlapped substantially between the subspecies, and the most extreme outlier belonged to the Kojima population, an insular population of *M. f. fuscata*. In the sex-combined analysis, there were no significant main effects of sex or subspecies. However, the sex–subspecies interaction effect was significant (0.33 [0.07, 0.59]) when excluding anomalous individuals, although this effect was not significant in the total sample (Tables 1 and S8).

**Figure 7.**
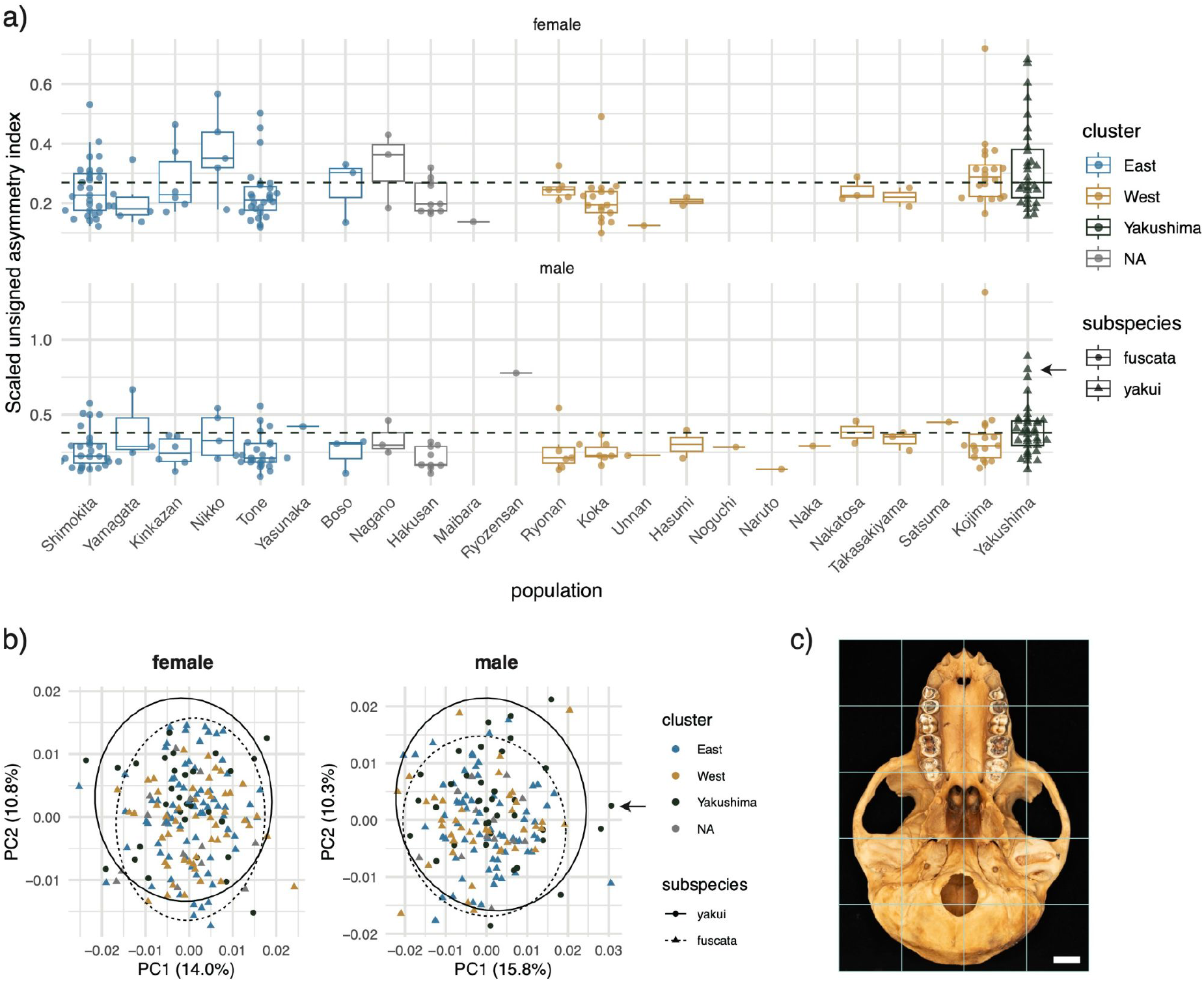
Fluctuating asymmetry as a measure of developmental instability. (a) Population differences in the scaled unsigned asymmetry index of cranial shape. Horizontal dashed lines indicate the median for *M. f. yakui*. (b) Scatter plots of the first two PCs of cranial asymmetric shape components for females (left) and males (right). (c) Example of a *M. f. yakui* cranium (PRI-7999) exhibiting a high unsigned asymmetry index. A grid has been overlaid to highlight the deformation. The neurocranium is distorted to the right relative to the face and palate (leftward in occlusal view). Scale bar indicates centimeters. Arrows in panels (a) and (b) identify this specimen.

**Table 1.** Bayesian regression estimates for inter-subspecies differences in the scaled unsigned asymmetry index. The models included population as a random effect.

|  |  | Estimate | Est. Error | Lower 95% CrI | Upper 95% CrI | Rhat | Bulk ESS | Tail ESS |
| --- | --- | --- | --- | --- | --- | --- | --- | --- |
| *Females* |  |  |  |  |  |  |  |  |
| Random effects | Population SD (intercept) | 0.155 | 0.081 | 0.033 | 0.344 | 1.004 | 1439.3 | 1659.9 |
| Fixed effects | Intercept | -1.436 | 0.070 | -1.566 | -1.288 | 1.004 | 2698.1 | 2487.5 |
|  | Subspecies (yakui) | 0.223 | 0.190 | -0.161 | 0.616 | 1.001 | 2909.2 | 2793.6 |
| Residuals | Sigma | 0.358 | 0.021 | 0.319 | 0.402 | 1.000 | 5608.2 | 5484.8 |
| *Males* |  |  |  |  |  |  |  |  |
| Random effects | Population SD (intercept) | 0.116 | 0.088 | 0.006 | 0.334 | 1.003 | 1812.5 | 3152.1 |
| Fixed effects | Intercept | -1.400 | 0.071 | -1.532 | -1.253 | 1.000 | 4178.5 | 3514.9 |
|  | Subspecies (yakui) | 0.404 | 0.171 | 0.054 | 0.750 | 1.001 | 3211.7 | 2752.3 |
| Residuals | Sigma | 0.438 | 0.028 | 0.388 | 0.498 | 1.000 | 7164.5 | 5424.6 |
| *Combined* |  |  |  |  |  |  |  |  |
| Random effects | Population SD (intercept) | 0.162 | 0.073 | 0.054 | 0.338 | 1.001 | 1582.2 | 1804.7 |
| Fixed effects | Intercept | -1.440 | 0.072 | -1.581 | -1.292 | 1.001 | 2852.7 | 3132.4 |
|  | Sex (male) | 0.036 | 0.053 | -0.067 | 0.143 | 1.001 | 7161.3 | 5461.2 |
|  | Subspecies (yakui) | 0.230 | 0.200 | -0.172 | 0.632 | 1.001 | 3696.1 | 3557.0 |
|  | Sex:Subspecies | 0.179 | 0.109 | -0.040 | 0.387 | 1.001 | 6784.8 | 6400.4 |
| Residuals | Sigma | 0.392 | 0.016 | 0.361 | 0.426 | 1.001 | 8874.3 | 5943.5 |

### Test of neutrality of morphological diversification

The beta test showed that the slope of the regression of log between-population variance against log within-population variance did not deviate significantly from one for either females (*F*_1,20_ = 0.09, *p* = 0.772) or males (*F*_1,21_ = 4.20, *p* = 0.053) (Fig. 8a). This indicated that the null hypothesis of neutrality could not be rejected for between-population morphological diversification in *M. fuscata*.

**Figure 8.**
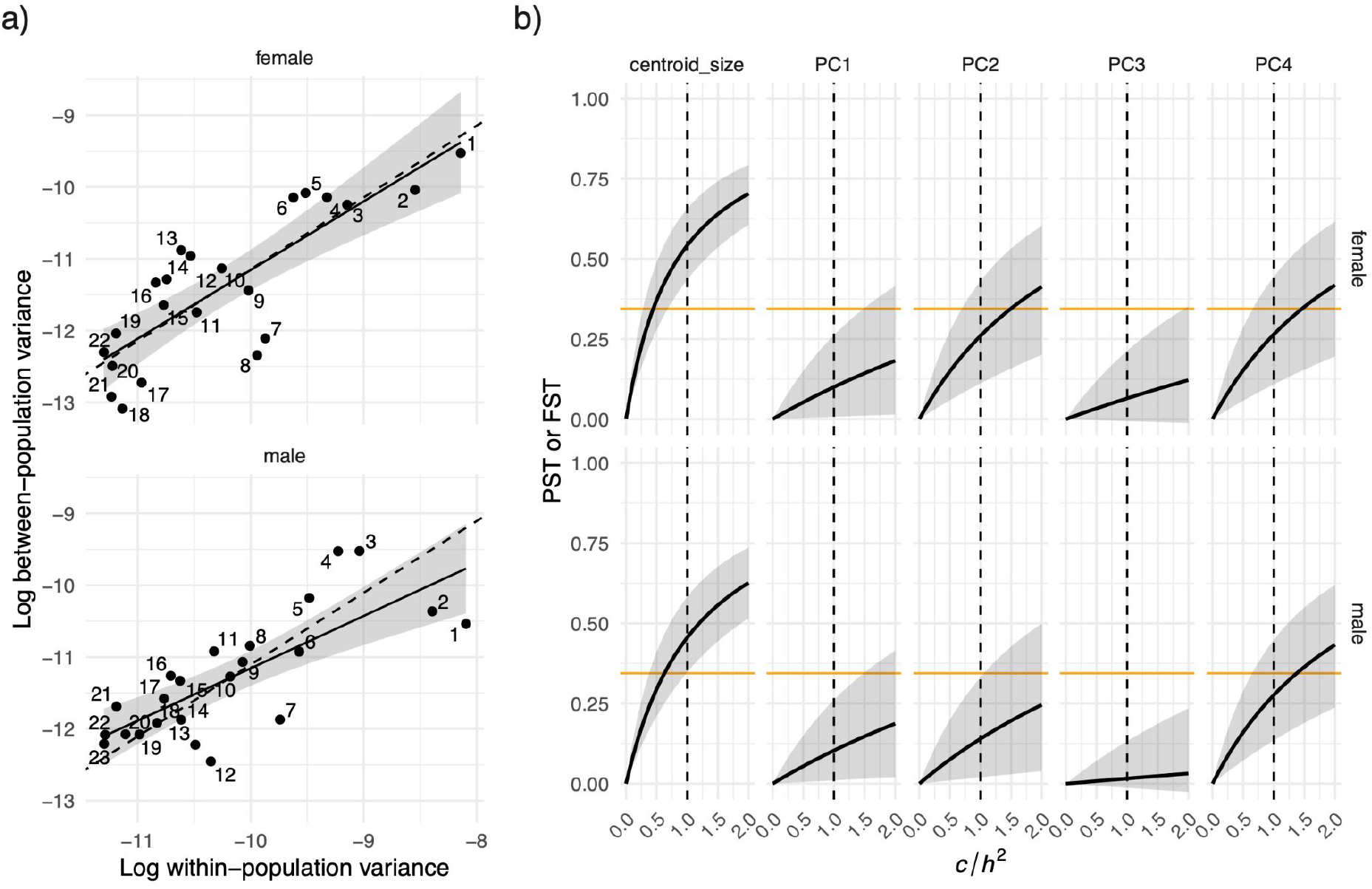
Neutrality tests for cranial diversity in *M. fuscata*. (a) Beta test comparing within-group and between-group variances. Solid lines and gray ribbons represent the regression and its 95% confidence intervals, respectively, while dashed lines indicate isometry. (b) *P_ST_* − *F_ST_* comparison between *M. f. yakui* and the West cluster (*M. f. fuscata*). The solid line represents *P_ST_* with 95% confidence intervals (gray ribbons). The horizontal orange line indicates the *F_ST_*. The vertical dotted line marks the null assumption (*c* = *h*^2^)

The multivariate drift test indicated that the observed magnitudes of divergence of *M. f. yakui* from *M. f. fuscata* were significantly smaller than the ranges expected under genetic drift. In females, the observed magnitude (95% CI: 0.022–0.032) fell below the simulated range (0.034–0.082). Similarly, in males, the observed magnitude (95% CI: 0.021–0.030) was smaller than the simulated range (0.035–0.085).

The *P_ST_* - *F_ST_* comparisons revealed that, for centroid size, *P_ST_* was significantly higher than *F_ST_* when assuming *c/h*^2^≥0.7 for females and *c/h*^2^≥0.1 for males (Fig. 8b). For the first four PCs, *P_ST_* did not significantly exceed *F_ST_* at any *c/h*^2^ values. Instead, the *P_ST_* values for PC1 and PC3 were significantly lower than *F_ST_* across most of the range of *c/h*^2^ values. Consequently, the size difference between *M. f. yakui* and the West clade (*M. f. fuscata*) indicated directional selection under certain *c/h*^2^ assumptions, whereas cranial shape differences were consistent with neutral processes or even indicated stabilizing selection, supporting the results of the multivariate drift test.

## DISCUSSION

### The origin and phylogenetic history of M. fuscata, with a particular focus on M. f. yakui

The maximum likelihood tree indicated that the *M. fuscata* lineage diverged from the ancestral lineage leading to *M. cyclopis*, Chinese *M. mulatta*, and Indian *M. mulatta*. This branching pattern corroborates the results of previous nuclear genome-based studies (Ito et al. 2021; Zhou et al. 2024; Higashino et al. 2025). Excess allele sharing was observed between Chinese *M. mulatta* and *M. fuscata*, and this sharing was more pronounced on the non-PAR X chromosome than on the autosomes. Consistent with the hypothesis initially proposed by Osada et al. (2021), our *f*-branch results support the scenario that male-biased gene introgression from the ancestral Chinese *M. mulatta* population into a population that branched off from the ancestral *M. fuscata* lineage caused nuclear swamping, leading to the formation of the current Chinese *M. mulatta* population. Furthermore, excess allele sharing was observed between *M. cyclopis* and *M. fuscata*, suggesting gene introgression between these two lineages before they migrated to Taiwan and the Japanese archipelago, respectively.

The migration history of *M. fuscata* ancestors to the Japanese archipelago remains a topic of debate, with some studies proposing a single migration wave (Kawamoto et al. 2007) and the others suggesting two or more waves (Nozawa et al. 1991; Hashimoto-Gotoh et al. 2020). Had a second migration event occurred, *M. fuscata* populations would likely exhibit heterogeneous genetic similarity to, and uneven excess allele sharing with, their continental relatives. However, the *f*-branch, *f*_3_, and *f*_4_ statistics revealed that both genetic similarity and excess allele sharing were uniform across *M. fuscata* populations. Furthermore, qpWave analysis did not support models with two or more migration waves. Collectively, these results support the hypothesis that the ancestors of *M. fuscata* migrated to the Japanese archipelago in a single wave.

The timing of migration to the Japanese archipelago is estimated at 430 or 640 ka (Marine Isotope Stage [MIS] 12 or 16) based on fossil records and paleo-sea-level estimations (Aimi 2002; Fooden and Aimi 2005). Our MSMC2 results identified a population decline shared by all *M. fuscata* populations around 400–500 ka. The *M. fuscata* common ancestors likely experienced a population bottleneck during the migration event in MIS 12.

The process of diversification within the Japanese archipelago has also been debated. The admixture analysis and PCA identified three major clusters: West (*M. f. fuscata*), East (*M. f. fuscata*), and Yakushima (*M. f. yakui*). The West–East boundary lies roughly between Koka in Shiga Prefecture (Ito et al. 2021) and Jigokudani in Nagano Prefecture. While Ito et al. (2021) attributed population divergence to isolation in glacial refugia [for the refugia hypothesis, see Otsuki et al. (2021)], Higashino et al. (2025) demonstrated that rCCR tends to increase during population declines. This suggests that glacial periods may have promoted admixture within refugia in at least some population pairs, rather than isolation.

Phylogenetic studies based on mitochondrial DNA (Kawamoto et al. 2007), blood protein variants (Nozawa et al. 1991), and reduced-representation sequencing of the nuclear genome (Ito et al. 2021) have challenged the sister relationship between *M. f. yakui* and *M. f. fuscata*. Our maximum likelihood tree also suggested that *M. f. yakui* was placed within the West clade of *M. f. fuscata*. However, our rCCR results suggested that the divergence of *M. f. yakui* (ca. 200–300 ka) predated the West–East divergence within *M. f. fuscata* (ca. 150–200 ka), supporting the sister relationship between the two subspecies. The estimated divergence time between *M. f. fuscata* and *M. f. yakui* (200–300 ka) is comparable to that estimated based on blood protein variants (178 ka) (Nozawa et al. 1977). The discrepancy in the branching pattern and divergence dates order can be attributed to the complex admixture history between the populations.

The NeighborNet graph shows high reticulation at the base of the three major clusters, supporting gene flows among populations. Both *f*-branch and *f*_4_ statistics suggest multiple episodes of sex-biased gene flow across these major clusters. TreeMix and admixture graph analyses identified two major ancestry waves: one contributing to the formation of present-day East clade populations, and the other shaping the extant *M. f. yakui* population. The ancestor of the West clade populations likely diverged from the *M. f. yakui* lineage, but subsequently underwent varying degrees of admixture from East clade lineages. The extent of this admixture aligns with geographic expectations, as the Kyushu populations were less admixed than the Shikoku–Kinki populations. Discrepancies in *f*-branch and *f*_4_ statistics between the autosomes and the non-PAR X chromosome suggest that male-biased migration drove the nuclear swamping.

The rCCR results indicate that gene flow between the *M. f. yakui* lineage and West clade lineages ceased or was severely restricted around 100 ka, even though a land bridge likely formed during the Last Glacial Maximum (LGM; MIS 2). The *M. f. yakui* ancestors likely experienced gene flow with West clade populations before migrating to Yakushima Island via the land bridge formed during MIS 6 (130–200 ka). Subsequently, the *M. f. yakui* population became isolated on the island.

### Long-term isolation, small population size, and inbreeding in Yakushima Island

The heterozygosity of wild *M. f. yakui* averages 0.0295% (0.0223–0.0318%). This is roughly one-third of that observed in most *M. f. fuscata* individuals and is one of the lowest values among primates (Kuderna et al. 2023). Comparable levels of low heterozygosity are found in the golden snub-nosed monkey (*Rhinopithecus roxellana*; 0.041%), and among other mammals, in the mainland clouded leopard (*Neofelis nebulosa*; 0.0406%), the Sunda clouded leopard (*N. diardi*; 0.0154%), the snow leopard (*Panthera uncia*; 0.0430%) (Yuan et al. 2023), and the ‘Southern Resident’ killer whale population (*Orcinus orca*; 0.029%) (Kardos et al. 2023). *M. f. yakui* thus represents an intriguing case study regarding extremely low genetic diversity and its potential impact on fitness and viability.

Regarding the causes of this extremely low genetic diversity, MSMC2 results indicated that *M. f. yakui* has experienced a decrease in effective population size since diverging from the *M. f. fuscata* lineage around 200–300 ka and has been isolated since around 100 ka. Our MSMC2 and SMC++ results detected a minor bottleneck around 10–20 ka. This minor bottleneck is common to all *M. fuscata* populations and is likely due to global climatic cooling during the LGM. Therefore, the extremely low genetic diversity of *M. f. yakui* is likely attributable, at least in part, to over 100,000 years of isolation on the island.

Hayaishi and Kawamoto (2006) proposed that *M. f. yakui* underwent a severe population decline due to environmental destruction caused by the Kikai Caldera eruption approximately 7.3 ka. This hypothesis is plausible, as thick pyroclastic flow deposits from the eruption have been identified at various sites except for the southern part of Yakushima Island (Machida and Arai 1978; Geshi 2009), although the exact extent of their dispersal remains debated (Kobayashi and Higashi 2023). Our Fastsimcoal2 results suggest that the effective population size shrank to nearly one-third around 4.5 ka, a contraction scenario further supported by the lower frequency of singletons in the site frequency spectrum of *M. f. yakui*. This decrease may be a consequence of eruption-induced environmental disruption, and the slight recovery around 3–5 ka observed in our MSMC2 results may reflect subsequent ecological recovery, as previously suggested for the sika deer (*Cervus nippon*) on Yakushima Island (Agetsuma-Yanagihara et al. 2025). However, our MSMC2 and SMC++ results did not detect a population bottleneck specifically corresponding to this timing. Similarly, the Yakushima gentian (*Gentiana yakushimensis*) exhibits extremely low genetic diversity yet does not appear to have experienced a population bottleneck during this period (Ishii et al. 2022). Furthermore, the genetic diversity of sika deer on Yakushima Island is not particularly low compared to that of mainland populations across the Japanese archipelago (Terada and Saitoh 2018; Liu et al. 2025). Interpreting these contrasting results remains challenging, as site-frequency-spectrum-based approaches have limited precision in inferring recent demographic events when sample sizes are small (Robinson et al. 2014). Therefore, although the pyroclastic flow likely covered a large area of the island, further studies with larger sample sizes are needed to evaluate the extent to which it impacted the population size of specific taxa.

The proportion of ROH (≥ 100 kb) is much higher in *M. f. yakui* than in *M. f. fuscata* populations. However, when considering only long ROHs (≥ 2 Mb), there is no significant difference between the two subspecies. This suggests that *M. f. yakui* has not specifically experienced elevated levels of inbreeding in the past few hundred years. Rather, the ROHs observed in *M. f. yakui* are likely the result of demographic events in the distant past. A similar pattern has been observed in Ethiopian wolves (*Canis simensis*), which are thought to have experienced long-term small population size and extensive genetic drift rather than recent inbreeding (Mooney et al. 2023). Historical events that could have contributed to the genomic structure of *M. f. yakui* include environmental disruption from the Kikai Caldera eruption, global climatic cooling during the LGM, and long-term isolation on the island. However, disentangling the relative contributions of these factors remains challenging.

### Genetic load in M. f. yakui

In smaller populations, genetic drift is more likely to outweigh purifying selection, causing the accumulation of deleterious alleles. Supporting this theoretical prediction, the nonsynonymous-to-synonymous ratio for missense mutations is much higher in *M. f. yakui* than in *M. f. fuscata* populations. The *R_XY_* results also revealed that LoF and missense derived alleles accumulated to a greater extent in *M. f. yakui* than in the West and East clades. These findings suggest that *M. f. yakui* experiences a higher drift load than *M. f. fuscata*.

Inbreeding converts masked load to realized load, which manifests as increased inbreeding depression and allows strongly deleterious alleles to be purged (Dussex et al. 2023a). As predicted by this theoretical expectation, the number of strongly deleterious (LoF) alleles tends to decrease in individuals with higher inbreeding coefficients within *M. f. fuscata*. However, *M. f. yakui* deviates from this trend, carrying a larger number of LoF alleles despite having much higher inbreeding coefficients. Such a non-linear (quadratic) relationship is also observed in montane red foxes (*Vulpes vulpes*) in North America (Quinn et al. 2024) and likely reflects the antagonism of selection and drift. Although purifying selection efficiently purges strongly deleterious alleles from smaller populations to a certain extent, the effect of selection is overwhelmed by genetic drift in *M. f. yakui*.

For LoF variants, traces of purifying selection are nevertheless observed in *M. f. yakui*. In the site frequency spectrum, deleterious mutations with stronger effects, particularly LoF variants, are more frequently observed as singletons, while they are less frequently observed at allele counts of two or more. This skewness is more marked in *M. f. yakui* than in the West and East clades of *M. f. fuscata*. Although the total load (i.e., the number of deleterious alleles) is greater in *M. f. yakui* than in *M. f. fuscata* populations, the difference is smaller for LoF variants than for missense variants. These results suggest that stronger purifying selection acted on LoF variants than on other mutation categories in *M. f. yakui*.

Genetic load in terms of lethal equivalents is shaped by the complex interplay between demographic history and selection coefficients (Kardos et al. 2021; Bertorelle et al. 2022; Dussex et al. 2023a). Some empirical studies have found efficient purging of strongly deleterious alleles, but not moderately deleterious ones, in populations that experienced historical bottlenecks (Xue et al. 2015; Robinson et al. 2018; Grossen et al. 2020; Khan et al. 2021; Dussex et al. 2023b), while others have not (Wang et al. 2021; Taylor et al. 2024). In *M. f. yakui*, both LoF and missense alleles accumulated to a greater extent than in the West and East clades; this accumulation pattern broadly echoes the latter findings. Theory predicts that purifying selection outweighs genetic drift when |s|≫1(2N_e_), where *s* is the selection coefficient and N_e_ is the effective population size (Robinson et al. 2023). It is reasonable to hypothesize that such severely deleterious variants have been at least partly purged in *M. f. yakui*; however, variants falling below this threshold |s|≤1(2N_e_) were likely fixed during a historical bottleneck and a period of long-term small population size and could not be removed by selection thereafter, resulting in drift load (Dussex et al. 2023a). Consequently, *M. f. yakui* carries, on average, a greater realized load and suffers from higher drift load than *M. f. fuscata* populations, while its potential load is comparable to or smaller than theirs.

### The potential morphological consequences of genetic load

We evaluated the fluctuating asymmetry of cranial shape as a measure of developmental instability. On average, the unsigned asymmetry index was higher in *M. f. yakui* than in *M. f. fuscata* for males, whereas no significant difference was found for females. This result remained consistent even when individuals with missing landmarks or qualitative anomalies were excluded. The sex–subspecies interaction effect was significant when the analysis was restricted to intact specimens.

Males may generally be more susceptible to genetic load because recessive deleterious alleles on the X chromosome are always exposed (hemizygous), whereas they are masked in females unless present in a homozygous state. However, there is no consensus as to whether males exhibit a greater degree of fluctuating asymmetry than females (Graham and Özener 2016). Our data also showed that there was no main effect of sex on the degree of fluctuating asymmetry, although the sex-subspecies interaction was significant when restricted to intact specimens. In primates, males tend to grow faster around puberty than females (Wang et al. 2007; Turcotte et al. 2022), and cranial sexual dimorphism manifests through ontogenetic scaling and divergent growth trajectories during later developmental periods (O’Higgins and Collard 2002; Cobb and O’Higgins 2007). These male-specific developmental demands may underlie male vulnerability to genetic load and higher susceptibility to developmental instability compared to females. Deleterious mutations that disrupt homeostatic development during the male-specific pubertal growth spurt or ontogenetic pathways may have accumulated in *M. f. yakui*, disproportionately affecting males.

An alternative hypothesis is that males are more sensitive to environmental stress than females. In macaques, intense male–male competition for mates likely exerts sexual selection pressure that drives rapid growth during puberty. On the island, where resources are limited, males may be forced to prioritize this rapid growth at the expense of developmental homeostasis, whereas females can prioritize stability. Thus, the male-specific increase in fluctuating asymmetry in *M. f. yakui* may reflect environmental stress caused by limited insular resources, rather than solely genetic load.

Regarding comparative studies, McGrath et al. (2022) reported that the facial crania of mountain gorillas, which exhibit extremely low genetic diversity and high inbreeding, possess a greater magnitude of fluctuating asymmetry than other gorilla subspecies. Island human populations also show higher levels of fluctuating asymmetry in the dental arch than reference populations (Schaefer et al. 2006). In other mammals and birds, populations on islands or with lower genetic diversity have been reported to show elevated levels of fluctuating asymmetry in skulls compared to reference populations (Hutchison and Cheverud 1992; Karvonen et al. 2003; White and Searle 2008; Lovatt and Hoelzel 2011). However, Caccavo et al. (2021) revealed that environmental stress, not genetic diversity, was related to the degree of fluctuating asymmetry in rodents. Although significant effects of inbreeding on fluctuating asymmetry have been reported for the wing veins of flies (*Drosophila melanogaster*) (Carter et al. 2009), there is no general consensus on whether inbreeding consistently increases fluctuating asymmetry (Waldmann 1999; Kruuk et al. 2003) or not (Fowler and Whitlock 1994; Lacy and Horner 1996; Rao et al. 2002; Kruuk et al. 2003). Furthermore, to the best of our knowledge, a male-specific effect such as that observed in the present study has not been reported previously.

The degree of cranial asymmetry observed in *M. f. yakui* appears considerably milder than the extreme examples observed in mountain gorillas (McGrath et al. 2022), and comparable levels of asymmetry are occasionally seen in some *M. f. fuscata* populations. Although our study suggests that genetic load increases fluctuating asymmetry—a measure of developmental instability—specifically in males, the generality of this finding warrants further investigation. If such an effect exists, it is likely to be relatively small, and the extent to which it is driven by genetic load versus environmental stress remains to be elucidated.

### Evolutionary drivers of morphological differentiation

*M. fuscata* inhabits diverse environments ranging from subtropical evergreen broad-leaved forests on Yakushima Island to cool-temperate deciduous forests on the Shimokita Peninsula. Significant inter-population variation in skull morphology has been reported in *M. fuscata*. Previous studies have suggested that cranial shape (Buck et al. 2018, 2019) and nasal cavity size (Rae et al. 2003) covary with ecogeographical variables such as temperature and precipitation, implying ecological adaptation. However, our beta test failed to reject the null hypothesis that between-population cranial shape differences resulted from genetic drift.

We subsequently focused on the cranial shape differentiation of *M. f. yakui* from *M. f. fuscata*. Our data support previous reports of cranial shape differences between the two subspecies (Yano et al. 2020). *M. f. yakui* is characterized by a generally narrower cranium—with the exception of the zygomatic arch—and exhibits a more peramorphic shape relative to *M. f. fuscata*, despite its smaller size. Neither our multivariate drift test nor our *P_ST_*- *F_ST_* comparison rejects the null hypothesis of neutrality; instead, they suggested stabilizing selection. This is consistent with the well-known phenomenon that quantitative traits often show substantial heritable variation despite being subject to stabilizing selection (Johnson and Barton 2005; Haller and Hendry 2014; Ponce de León et al. 2018). Cranial shape may have diversified stochastically but within strong constraints that pull the populations back toward a common adaptive peak.

Note that failure to reject the null hypothesis does not necessarily mean that directional selection did not operate during morphological diversification. Another limitation of our analyses is the relatively small sample size used to calculate within-population V/CV matrices, which may bias the estimation of ***G***. Despite these limitations, our results suggest that stochastic processes operating under evolutionary constraints should be considered as a starting point for discussing the observed cranial shape diversity. Due to its long-term small population size, the distinctive cranial shape of *M. f. yakui* may have evolved through genetic drift under stabilizing selection.

Cranial size, a proxy for body size, is significantly smaller in *M. f. yakui* than in *M. f. fuscata*. The *P_ST_* - *F_ST_* comparison rejected neutrality when assuming **c/h*^2^*≥0.7for females and *c/h*^2^≥1.0 for males. Here, the assumption *c/h*^2^ ≈ 1 implies that the proportion of phenotypic variance due to additive genetic effects is comparable for between-population variance and within-population variance (Brommer 2011). Despite the long-term small population size, directional selection could have acted on body size on Yakushima Island, as predicted by the island rule. Sexual selection may have opposing effects on body size in males, which might have obscured the signature of directional selection.

## CONCLUSION

This study examined the genomic and morphological consequences of long-term isolation and small population size, using *M. f. yakui* as a case study. Population genetic analyses revealed that *M. f. yakui* diverged from *M. f. fuscata* and has remained isolated on Yakushima Island for over 100,000 years, maintaining a small effective population size of several thousand individuals. This demographic history has resulted in extremely low genetic diversity—approximately one-third of that observed in *M. f. fuscata*—with more than 70% of the genome covered by ROH ≥ 100 kb. Due to the pervasive effects of long-term genetic drift, deleterious mutations were prone to fixation and consequently were not efficiently purged. The resulting increase in realized genetic load likely caused developmental instability, manifesting as elevated levels of fluctuating asymmetry in cranial shape, particularly in males. Environmental stress on the island may alternatively account for this male-specific increase in fluctuating asymmetry. Despite the predominance of genetic drift, directional selection might have driven the reduction in body size in *M. f. yakui*, as predicted by the island rule. In contrast, cranial shape differentiation is consistent with a scenario of evolution driven by genetic drift acting under the constraints of stabilizing selection.

## ACKNOWLEDGEMENTS

We would like to express our gratitude to Hideo Higashi, Toshihito Takagi, Hidetoshi Tamate, Yusuke Hamazaki, Arisa Ishimura, Nami Suzuki-Hashido, Mayumi Morimoto, Yuzuru Hamada, Mikiko Tanaka, and Hiroo Imai for their help with sample preparation; Kosei Izawa, Tetsu Hayashi, Terutake Hayashi, Emiko Ashida, Hironori Seino, and all others who contributed to preserving the skeletal specimens and helped our study; Makoto Shimada, Atsunori Higashino, and Naoki Osada for providing information on relevant samples or data; and the successive students and staff of the Department of Evolution and Phylogeny, Primate Research Institute, Kyoto University, for their efforts in collecting, preparing, and preserving the skeletal collections. We thank Oita City for permitting the use of the Takasakiyama samples and specimens; Chiba Prefecture for permitting the use of the Boso specimens; the Ministry of the Environment, the Yakushima Forest Ecosystem Conservation Center, the Kagoshima Prefectural Government, and the Cooperation Research Program of the Wildlife Research Center at Kyoto University for granting permission to conduct research in Yakushima; and the McDonnell Genome Institute at Washington University for making the macaque genome assembly data available. We used GitHub Copilot, ChatGPT, and Gemini for coding assistance, as well as DeepL Write and Gemini for English proofreading and critical pre-submission feedback. We carefully verified and edited the AI-generated suggestions to ensure accuracy; full responsibility for the analysis scripts and manuscript rests with the authors.

## DATA AVAILABILITY

Raw sequencing reads have been deposited in the DDBJ Sequence Read Archive under BioProject accession number PRJDB38086. Additional data and analysis scripts are available on Dryad (https://doi.org/10.5061/dryad.brv15dvrn) and Zenodo (https://doi.org/10.5281/zenodo.22636544), respectively.

## AUTHOR CONTRIBUTIONS

T.I., T.Ha., and Y.Ka. conceived the project. T.I., Y.Ku., G.H., A.K., T.N., S.A., and T.Ho. conducted a field survey on Yakushima Island. S.Y., T.A., T.O., S.H., and Y.Ka. prepared the samples and specimens. T.I. and H.W. performed the molecular experiments and CT scanning. T.I. analyzed the data and drafted the manuscript, with input from the other authors. All authors approved the final manuscript.

