## Supplementary Figures for "Genomic and phenotypic consequences of long-term isolation and small population size in Yakushima macaques"

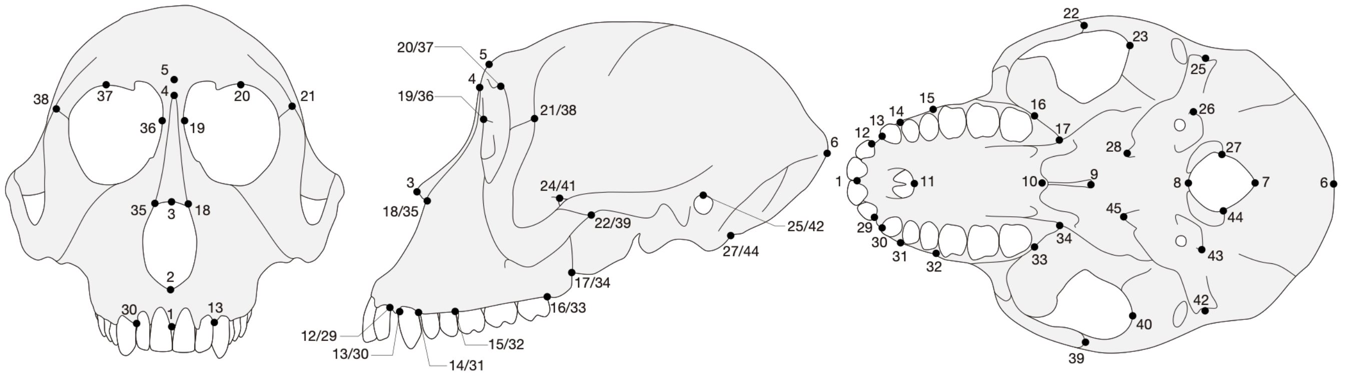


**Figure S1.** Cranial landmarks digitized for geometric morphometrics

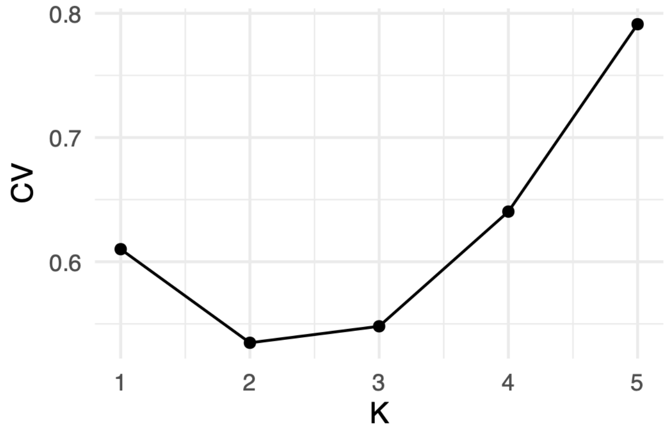


**Figure S2.** Cross-validation error in ADMIXTURE analysis across K values from 1 to 5.

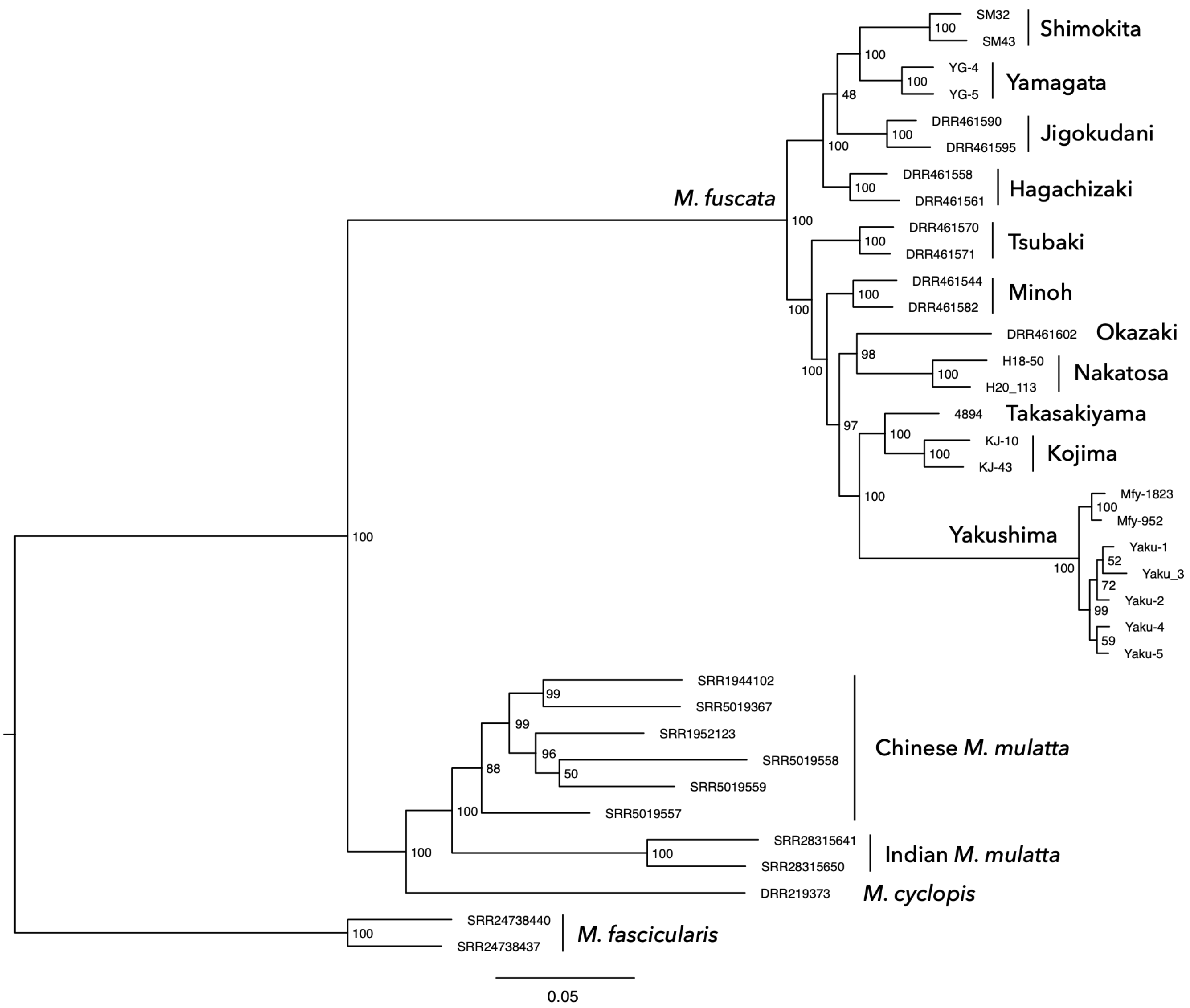


**Figure S3.** Maximum likelihood tree based on the autosomal SNVs. Bootstrap support values are shown at the nodes.

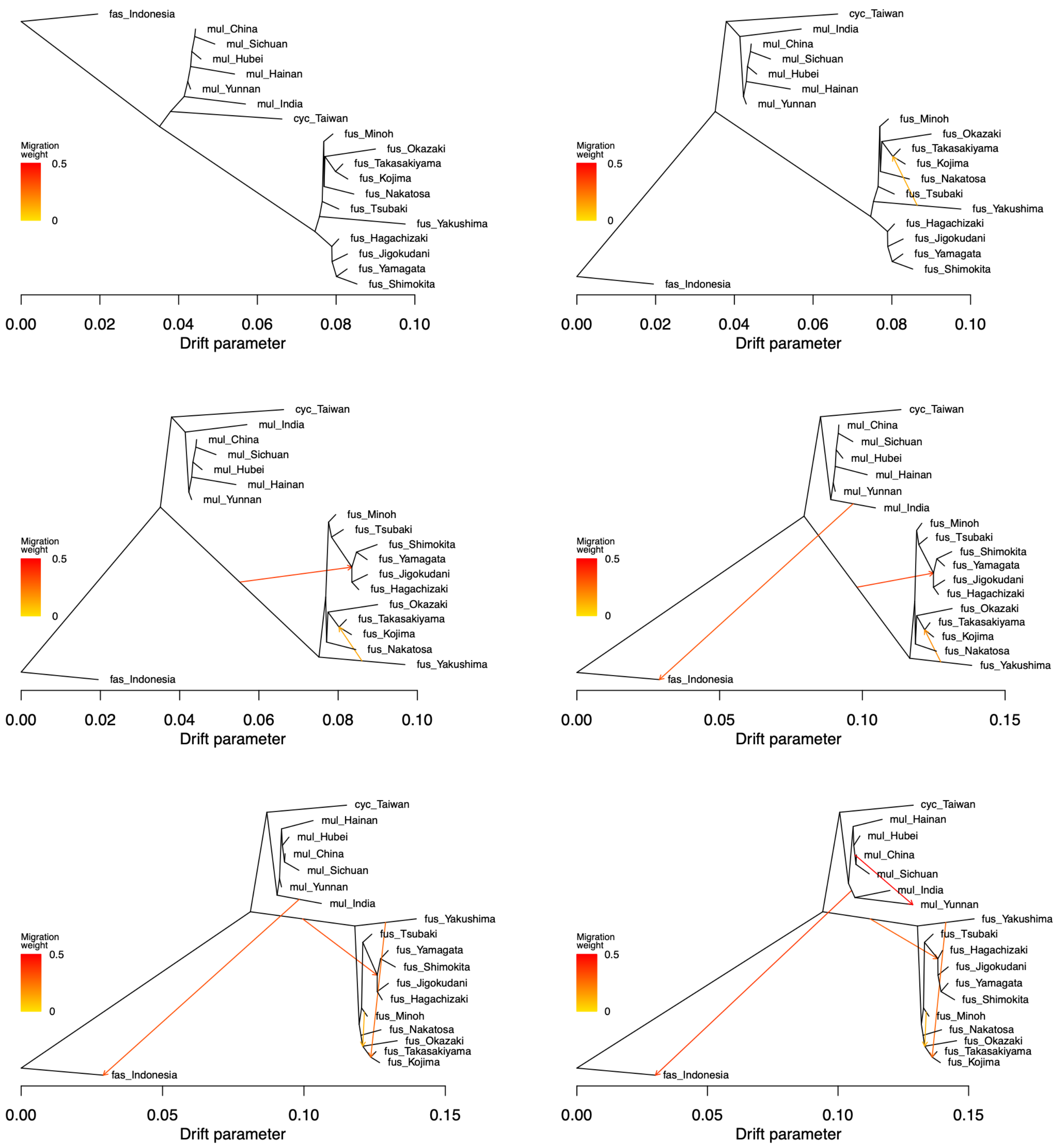


**Figure S4.** TreeMix results with 0 to 5 migration edges (from top left to bottom right). Abbreviations: fus, *M. fuscata*; cyc, *M. cyclopis*; mul, *M. mulatta*; and fas, *M. fascicularis*.

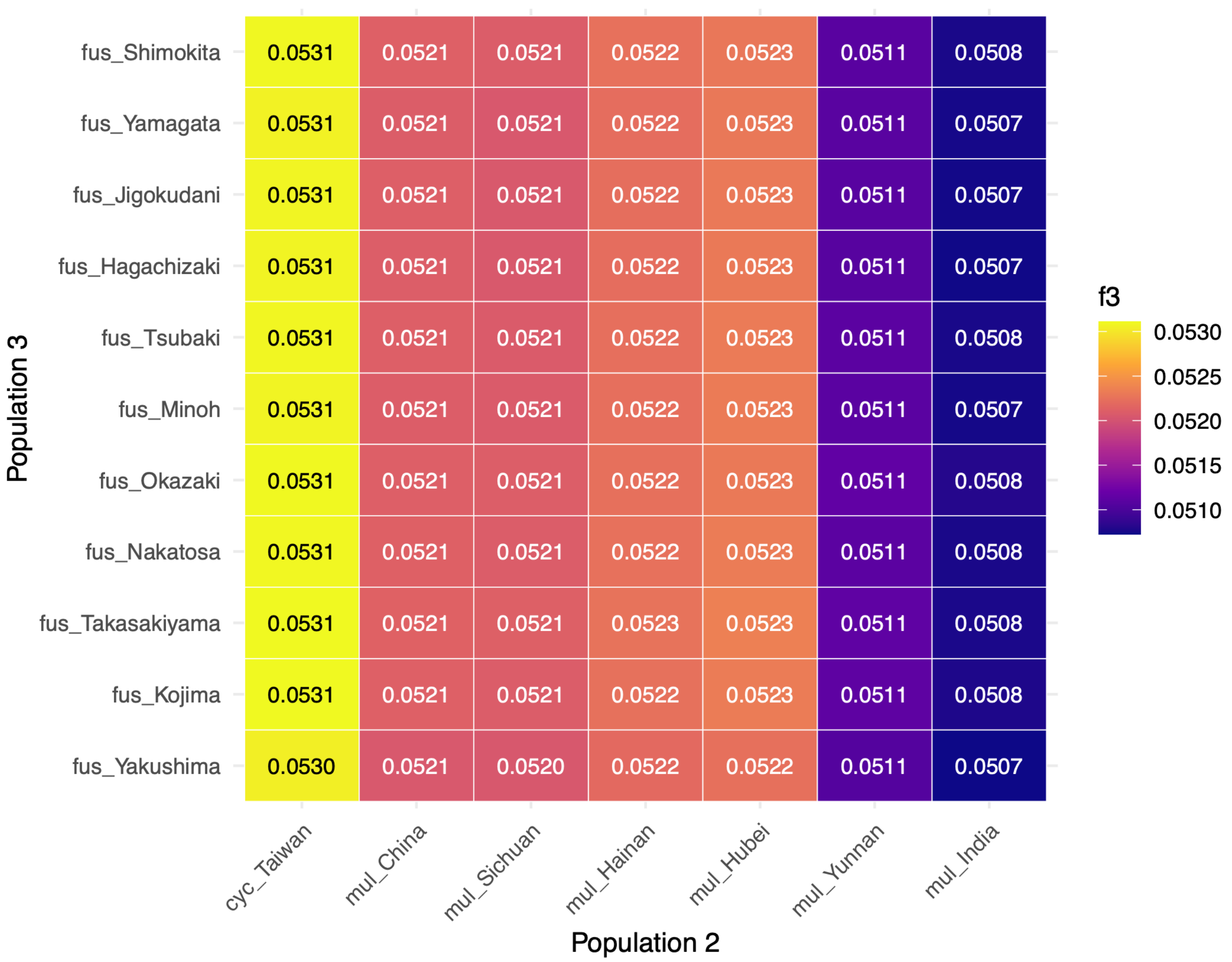


**Figure S5.** Autosomal outgroup $f_3$ statistics in the form of $f_3$ (fas_Indochina; P2, P3). Higher values indicate greater genetic affinity between P2 and P3. See the legend of Fig. S4 for abbreviations.

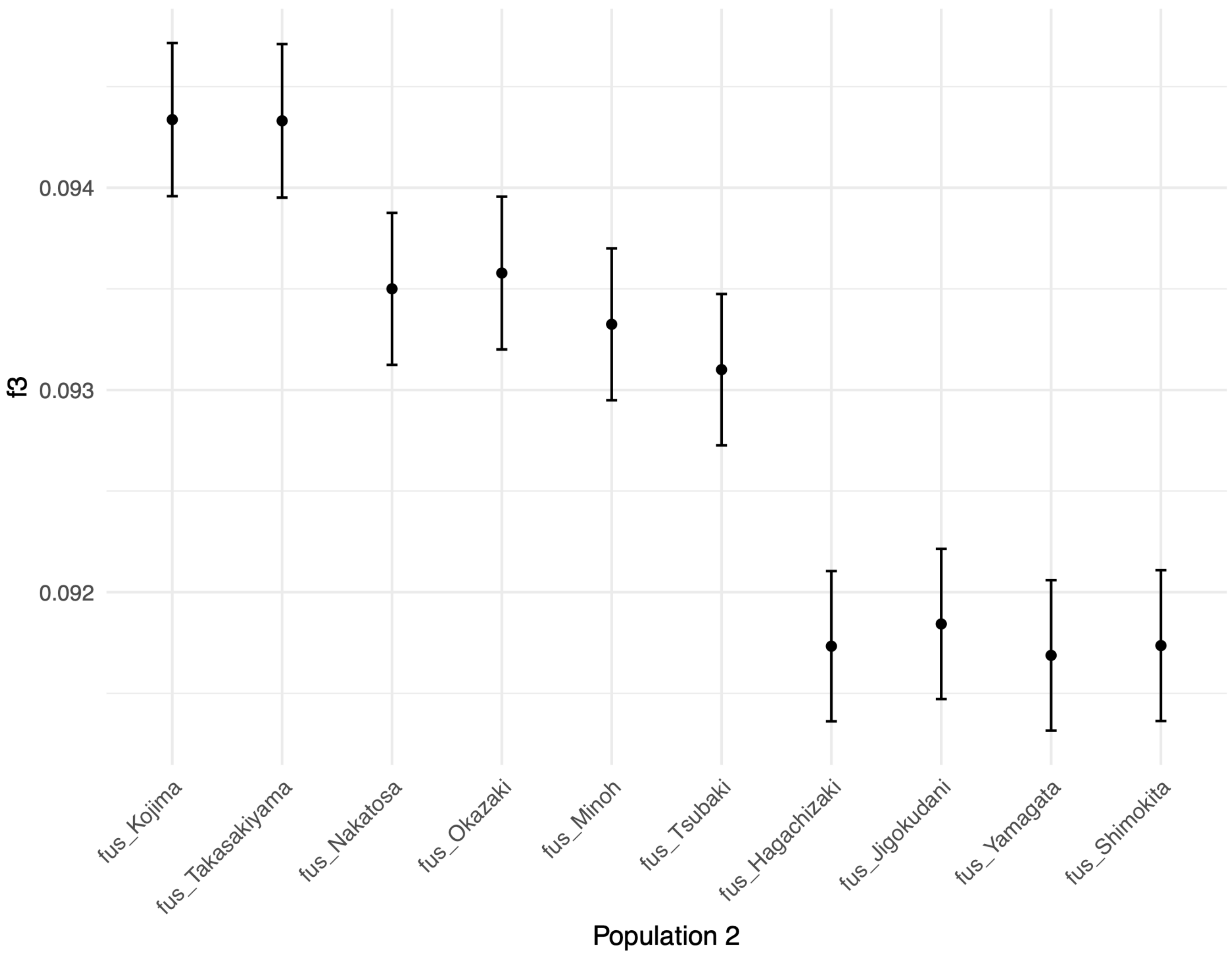


**Figure S6.** Autosomal outgroup $f_3$ statistics in the form of $f_3$ (fas_Indochina; P2, fus_Yakushima). Higher values indicate greater genetic affinity between P2 and fus_Yakushima. See the legend of Fig. S4 for abbreviations.

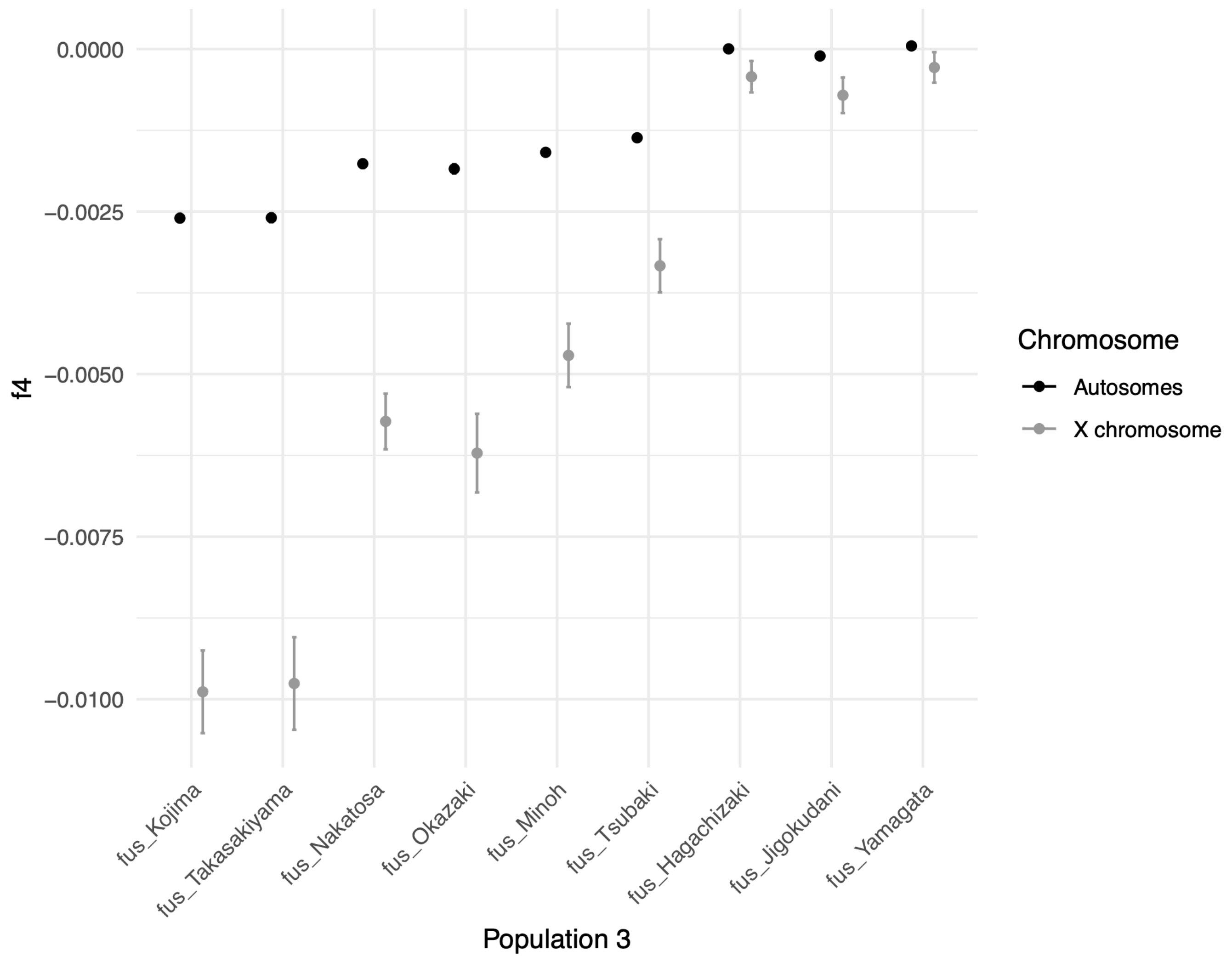


**Figure S7.** Autosomal and X-chromosomal $f_4$ statistics in the form $f_4$(fas_Indochina, fus_Yakushima; P3, fus_Shimokita). More negative values indicate a higher proportion of admixture between fus_Yakushima and P3. The discrepancy between autosomes and X chromosomes suggests sex-biased introgression. See the legend of Fig. S4 for abbreviations.

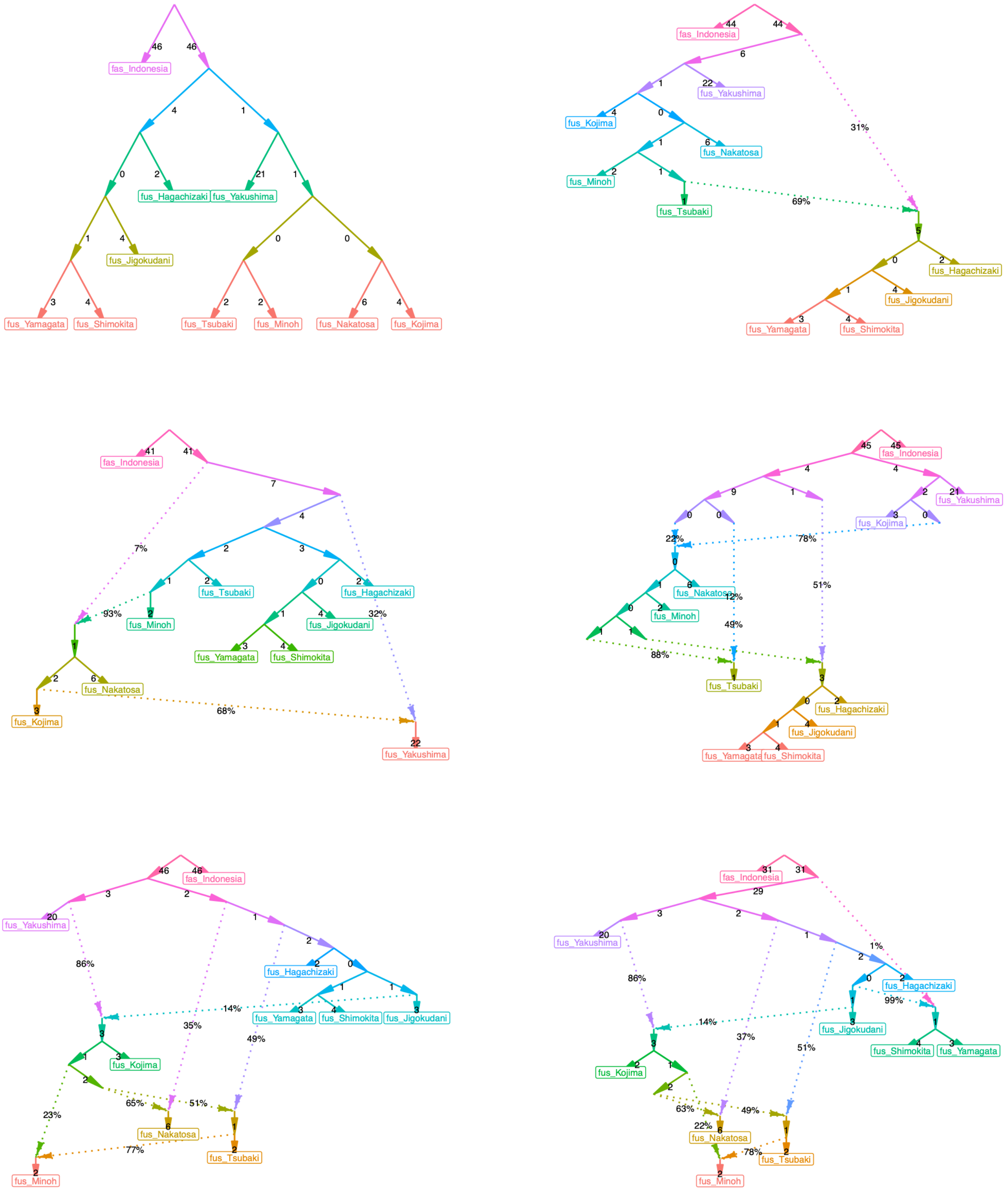


**Figure S8.** Admixture graphs for scenarios with 0 to 5 migration events (from top left to bottom right). See the legend of Fig. S4 for abbreviations.

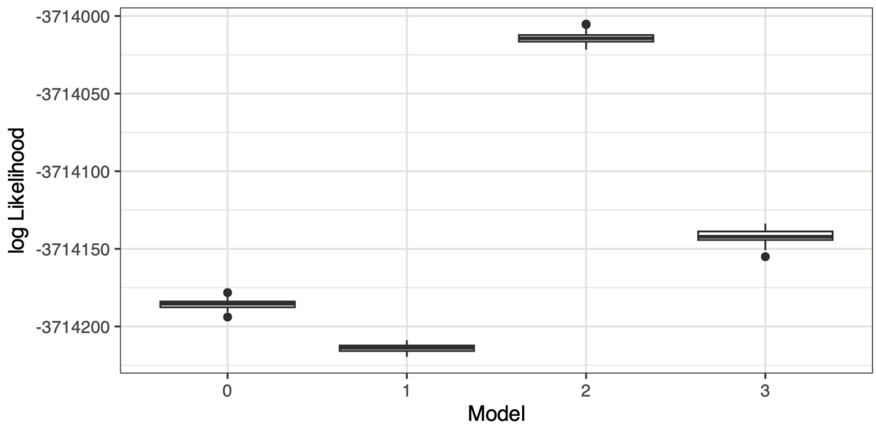


**Figure S9.** Likelihood distribution across models.

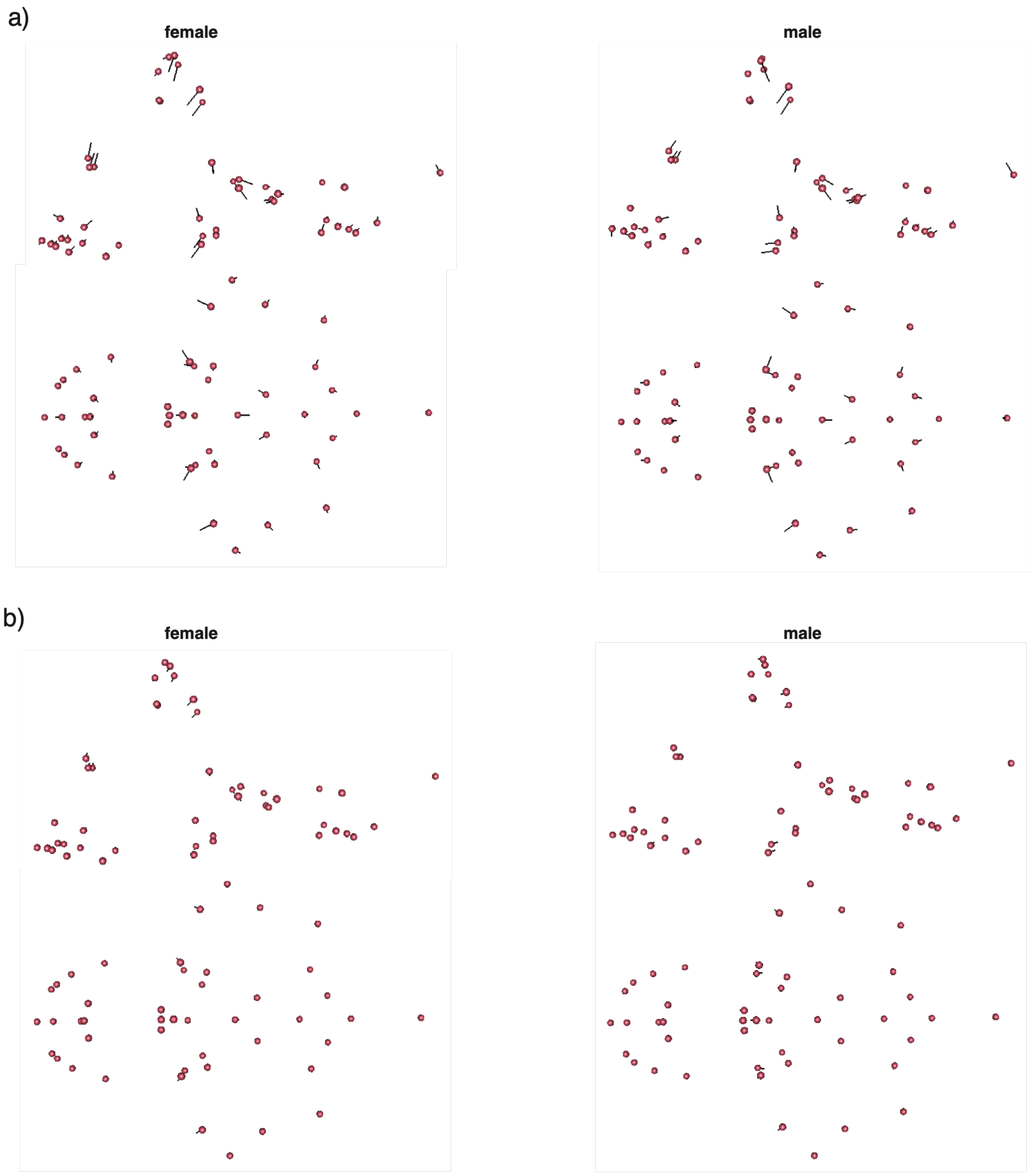


**Figure S10.** Vector-based visualizations of cranial symmetric shape variation in *M. fuscata*. (a) Inter-subspecies shape differences analyzed by between-group PCA. Points represent the reference (bgPC1 score −3 SD; *M. f. fuscata*) and vectors indicate the target (bgPC1 score +3 SD; *M. f. yakui*). (b) Allometric shape changes associated with regression scores. Points represent the reference (regression score −3 SD) and vectors indicate the target (regression score +3 SD). Shape changes visualized by deformed meshes are shown in Fig. 6.

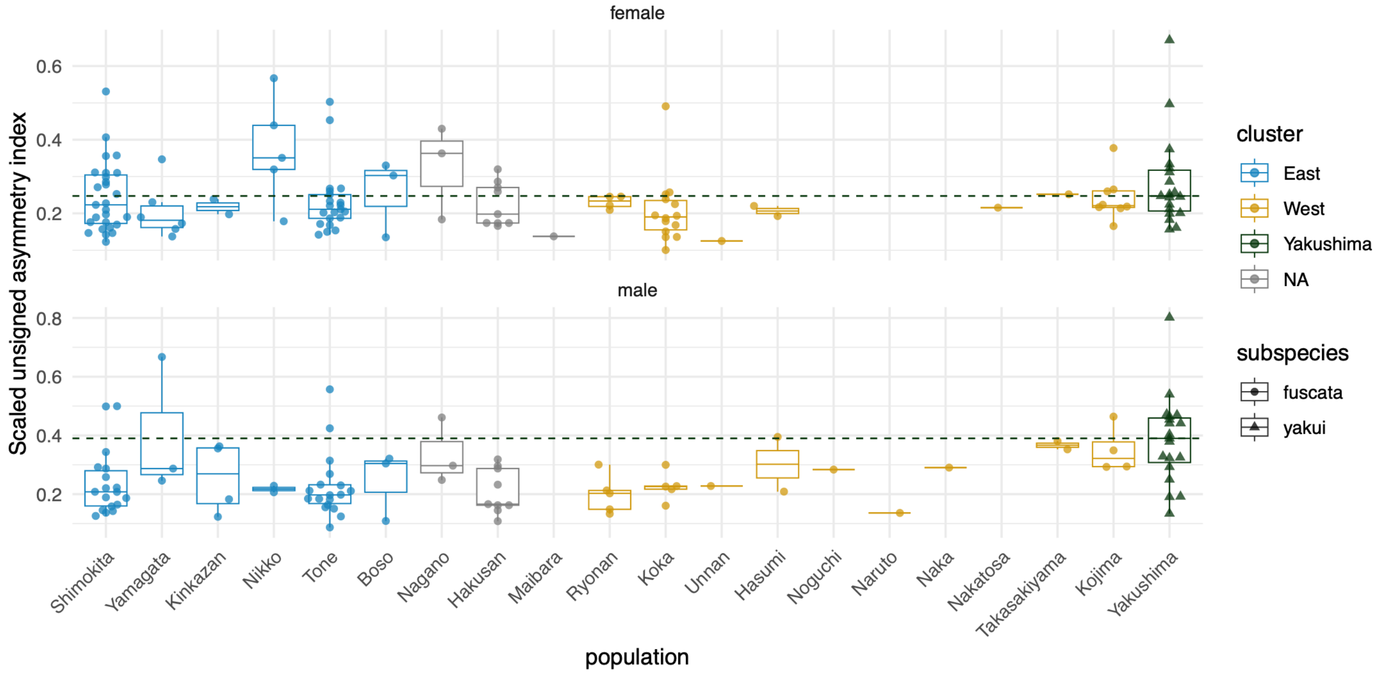


**Figure S11.** Population differences in the scaled unsigned asymmetry index of cranial shape. A total of 228 specimens with no missing landmarks or qualitative anomalies were included in this analysis. Horizontal dashed lines indicate the median for *M. f. yakui*.
